# Primary Human Macrophages Exhibit Functional CYP3A4 Activity with Marked Expression-Function Discordance and Cocaine-Induced Catalytic Activation

**DOI:** 10.64898/2026.10.21.760694

**Authors:** Amira S. Price, Stephen Knerler, Rodnie Colón Ortiz, Jareb J. Pérez-Caraballo, Alysha L. Ellison, Cory J. White, Jose Javier Rosado-Franco, Hannah N. Wilkins, Alicia Mercado, Kiara Mehta, Oluwatofunmi Oteju, Parker Stevenson-Li, Kimberly S. Williams, Peter J Gaskill, Dionna W. Williams

## Abstract

CYP3A4 is a phase I drug-metabolizing enzyme responsible for transformation of ∼50% of clinically relevant drugs. While extensively characterized in hepatocytes, its role in immune cells remains poorly understood. Here, using primary human monocyte-derived macrophages, we identify an understudied extrahepatic site of CYP3A4-mediated drug metabolism. Macrophages from 100 independent donors expressed CYP3A4 and its transcriptional regulators, Pregnane X Receptor (PXR) and Constitutive Androstane Receptor (CAR), with substantial interindividual variability. Importantly, CYP3A4 protein expression was accompanied by measurable catalytic activity in intact macrophages, yet CYP3A4 mRNA, protein abundance, and activity lacked concordance, demonstrating that expression alone may not predict function. *In vivo*, CYP3A4 was expressed in rhesus macaque Kupffer cells, further supporting macrophages as a potential pharmacologic entity. We next evaluated cocaine as a potential modulator of CYP3A4, due to its role in xenobiotic clearance. Cocaine had a dichotomous effect on expression, increasing CYP3A4 in some donors while producing no change or decreases in others, a phenotype associated with PXR, CAR, and the sigma-1 cocaine receptor. Strikingly, despite these divergent expression responses, cocaine increased CYP3A4 catalytic activity in all donors, reaching or exceeding levels induced by rifampicin, a clinically relevant CYP3A4 inducer. In rhesus macaques, cocaine also decreased antiretroviral therapy tissue concentrations, coincident with increased CYP3A4 and PXR. Together, these findings establish macrophages as a functionally active extrahepatic site of CYP3A4 metabolism and reveal a critical disconnect between CYP3A4 expression and catalytic function. Moreover, our findings identify interindividual variability and expression-activity discordance as important considerations for personalized medicine and understanding clinically relevant drug-drug interactions in the context of substance use.

**Significance statement:** CYP3A4 is best known for its role in hepatic drug metabolism, but its function in other cell types remains poorly understood. This work establishes macrophages as a functionally active extrahepatic site of CYP3A4-mediated drug metabolism, with marked interindividual variability and poor concordance between CYP3A4 expression and activity. Cocaine-enhanced CYP3A4 activity, together with altered antiretroviral tissue disposition *in vivo*, supports macrophage drug metabolism as a potential contributor to variability in therapeutic responses and substance use-associated drug-drug interactions.

**Visual abstract:** 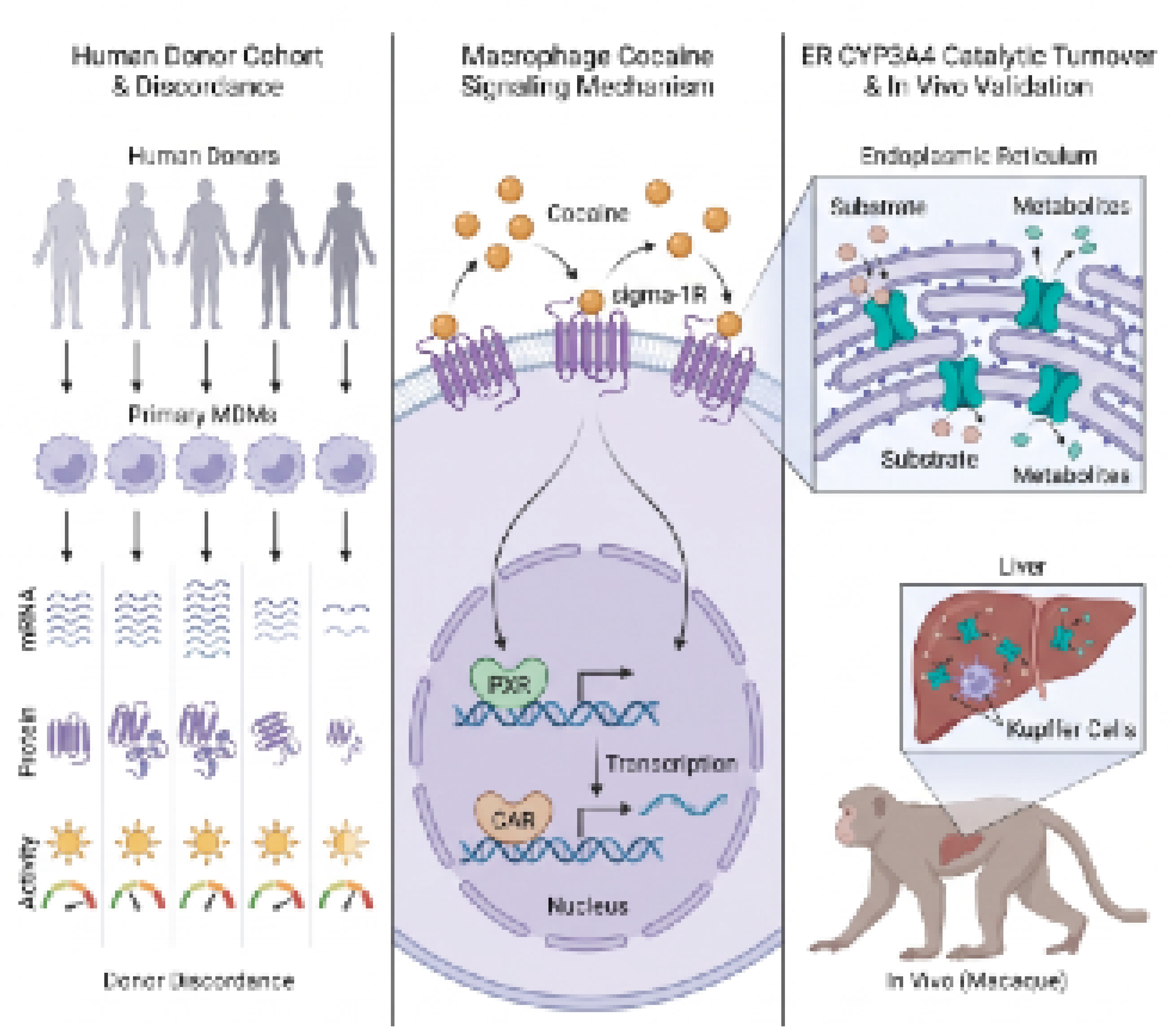

## Introduction

Cytochrome P450 3A4 (CYP3A4) is an enzyme of the cytochrome P450 family that metabolizes ∼50% of drugs in clinical use^1^. While CYP3A4 plays a critical role in phase I drug metabolism, it is highly promiscuous and participates in modification and clearance of xenobiotics, including addictive substances^2,3^. Additionally, as part of its broad substrate specificity, CYP3A4’s expression and activity are vulnerable to modulation by several of its ligands and common dietary components^4–6^. Through interactions with its transcriptional modulators, the nuclear receptors pregnane X-receptor (PXR) and constitutive androstane receptor (CAR), CYP3A4 expression and activity can be readily induced or inhibited. Through this PXR- and CAR-induced regulation, modulation of CYP3A4 activity can promote adverse drug:drug interactions of major clinical consequence as they impact drug clearance and the pharmacologic profile of CYP3A4 substrates^7–9^. Thus, a comprehensive understanding of the factors that influence CYP3A4 expression and activity is integral to navigating adverse drug:drug interactions and improving patient outcomes.

Liver and intestine homogenate analyses have been integral in understanding CYP3A4 activity and regulatory mechanisms. However, this approach does not permit investigation of cell type diversity within these compartments nor extrahepatic pharmacologic mechanisms in understudied cell types. This is important because understanding cell type-specific pharmacologic mechanisms can be predictive of relevant clinical outcomes. For example, evaluation of phase I drug metabolic mechanisms in immune cells is related to long-term therapeutic success for people with HIV taking antiretroviral therapies (ART)^10–15^. Similarly, studies using endoplasmic reticulum-enriched microsomes have also been integral in evaluating CYP3A4. However, this approach does not permit evaluation of CYP3A4 activity in intact, live cells, losing the ability to evaluate the intracellular nuances of CYP3A4-regulation, and results in low throughput readouts as microsomal preparation is time consuming and not well suited for evaluation of pharmacologic mechanisms among differing individuals. Moreover, methodologies that permit CYP3A4 investigation in intact, live cells from multiple donors represents an underutilized approach that can be important in identifying personalized medicine approaches.

To this end, our study focused on evaluating CYP3A4 expression and activity in macrophages – an important immune cell type expressed in all organs that contributes to both homeostasis and pathogenesis in a wide variety of disorders. As this has been an understudied cell type, we first set out to characterize CYP3A4 expression and activity using cells obtained from 100 individual donors. To our surprise, macrophages possessed the capability to participate in phase I reactions, as cells from all donors expressed CYP3A4 and its transcriptional regulators PXR and CAR. Indeed, CYP3A4 was not only expressed but functional, as determined by our activity assay that quantified the O-debenzylation of 7-benzyloxy-4-trifluoromethylcoumarin (BFC) to produce the fluorescent metabolite 7-hydroxy-4-trifluoromethylcoumarin (HFC). Additionally, macrophage CYP3A4 was subject to induction by canonical modulators dexamethasone and rifampicin. *In vivo*, macrophage CYP3A4 expression was confirmed using rhesus macaque Kupffer cells, extending the clinical relevance of our findings. To evaluate their capacity for contributing to adverse drug:drug interactions, macrophages were treated with cocaine. We determined that cocaine had variable effects on CYP3A4 in a donor-dependent fashion, where expression was increased, decreased, or remained unchanged depending on the individual. We term those for whom cocaine increased CYP3A4 expression “responders” and the remaining donors “non-responders”. We determined that the CYP3A4 expression “responsiveness” profile was unrelated to transcriptional network, CYP3A4 genotype, or demographic factors. Moreover, cocaine’s effects on CYP3A4 expression occurred in a cocaine-specific fashion, as cocaine metabolites or dopamine failed to replicate the CYP3A4 expression “responsiveness” profile. To this end, and unexpectedly, cocaine increased CYP3A4 activity for all evaluated donors, irrespective of their CYP3A4 expression “responsiveness” profile. Surprisingly, donors with the CYP3A4 “non-responder” expression profile had the greatest induction in CYP3A4 catalytic activity due to decreased basal levels of enzyme activity and increased endogenous levels of the cocaine receptor sigma-1R (σ1R). Finally, we performed proof of concept experiments using rhesus macaques that received ART in the presence and absence of cocaine, as substance use is inextricably linked to the HIV pandemic^16–19^ and our previous findings demonstrate that cocaine impacts ART disposition through PXR-mediated regulation^20^. We found that cocaine decreased tissue concentrations of two first-line ART therapies and their active metabolites, which coincided with cocaine-induced increases in CYP3A4 and PXR. We also confirmed the established premise that cocaine increases HIV infection of macrophages, reinforcing the pressing need to identify mechanisms to impact ART metabolism in macrophages, and therefore impact its antiviral efficacy, in people with HIV who also use cocaine and potentially other substances of abuse.

## Materials and Methods

### Human peripheral blood mononuclear cell (PBMC) isolation

Peripheral blood from healthy human volunteers was collected in a citrate-dextrose solution (Millipore Sigma, Burlington, MA) under protocols approved by the Emory University Institutional Review Board. Peripheral blood mononuclear cells (PBMC) were isolated by density-gradient centrifugation using Ficoll-Paque PLUS (Cytiva, Marlborough, MA) and 1X Dulbecco’s phosphate buffer saline (DPBS; Gibco, Waltham, MA). Red blood cells (RBC) were removed using 1X RBC lysis buffer prepared from a 10X stock solution in distilled water containing 1.5 M ammonium chloride (Sigma-Aldrich, St. Louis, MO), 0.10 M sodium bicarbonate (Sigma-Aldrich), and 0.01 M disodium ethylenediaminetetraacetic acid (EDTA; Sigma-Aldrich, St. Louis, MO). PBMC were resuspended in macrophage media made of Dulbecco’s Modified Eagle Medium (DMEM; Gibco) buffered with 15 mM HEPES (Quality Biological, Inc., Gaithersburg, MD) and supplemented with 10% fetal bovine serum (R&D Systems, Minneapolis, MN), 5% human serum AB (GeminiBio, Sacramento, CA), 1% penicillin–streptomycin (10,000 U/mL; Gibco), 2 mM L-glutamine (Gibco), and 1% M-CSF (R&D Systems).

### Monocyte derived macrophage (MDM) cell culture

PBMC were seeded in 10-cm tissue culture dishes (Fisher Scientific, Waltham, MA) and maintained at 37°C with 5% CO₂, with fresh macrophage media added on day three. After six days of culture, adherent cells demonstrating elongated or spindle-like morphology were considered mature MDM and used for downstream experiments.

### MDM cocaine, norcocaine, benzoylecgonine, and dopamine treatments

MDM were treated for 24 hours with cocaine hydrochloride (10 μM, NIDA Drug Supply Program, Research Triangle Park, NC), norcocaine (10 μM, NIDA Drug Supply Program), benzoylecgonine (10 μM, NIDA Drug Supply Program), or dopamine (10 μM, Millipore Sigma) and subsequently used in downstream experiments. Treatment with vehicle was used as a control.

### MDM HIV Infection

Mature MDM were exposed to 1 ng/mL HIV_ADA_ (NIH HIV Reagent Program, Division of AIDS, NIAID, NIH: Human Immunodeficiency Virus-1 ADA, ARP-416 contributed by Howard Gendelman) for 24 hours, the virus removed by adding fresh macrophage media, and the cells cultured for up to 9 additional days to facilitate viral replication. A portion of the cells remained uninfected as a negative control.

### Kupffer cell isolation

Rhesus macaque liver was obtained and a total of 50 grams of tissue from the periphery of the organ was removed for Kupffer cell isolation. Laennec’s capsule was removed and the tissue was gently washed with cold phosphate buffered saline (PBS; Gibco). The tissue was chopped using surgical (Cynamed, Gainesville, VA) and razor (WorkPro, Huntersville, NC) blades, removing and discarding vessels as necessary, until a consistency resembling a paste was reached. Cold PBS was added to the tissue paste to create a slurry capable of pipetting. The tissue slurry was further macerated by pipetting with a 50 mL serological pipette until tissue no longer clogged the pipette. The tissue slurry underwent enzymatic digestion with collagenase type 1 (final concentration 0.5 mg/mL per 5 mL of tissue; Worthington, Lakewood, NJ) and trypsin inhibitor (final concentration 0.01 mg/mL per 5 mL of tissue; Sigma-Aldrich, St. Louis, MO). The tissue slurry was incubated with digestion enzymes in a rotating incubator (Benchmark Scientific, Sayreville, NJ) for 1 hour, 37°C, rotation speed of 60 rpm, or until it appeared as a light pink homogenous liquid with little to no pieces of tissue remaining. Digestion lasted no longer than 1 hour to prevent over digestion and damage to cells. Additionally, after the first 25 minutes of enzymatic digestion, incubation was paused and the tissue was removed for additional maceration with a 10 mL serological pipette until tissue no longer clogged the pipette. After 1 hour, the digestion was halted by adding fetal bovine serum (FBS, final concentration 10%; VWR Radnor, PA) to the cell solution and a series of filter and wash steps was performed to remove residual tissue and digestion enzymes. The cell solution was filtered through a sterile 180 mm mesh (Homeemoh) followed by a 100 mm cell strainer (Corning), then pelleted (300 g, 5 minutes, 4°C) and washed three times with 4°C PBS. Note that the supernatant from the first wash was retained, centrifuged again to pellet additional cells, and this pellet washed three times with cold PBS as well. Following the washes, all cell pellets were combined and subjected to density gradient separation using 33% Percoll (700 g, 12 minutes, 4°C, no break; Cytiva, Marlborough, MA). The subsequent pellet was retained and resuspended in cold PBS, filtered through a 70 mm cell strainer (Corning), and subjected to RBC lysis for 10 minutes at 37°C. Cells were then pelleted (300 g, 5 minutes, 4°C), resuspended in cold PBS, placed on ice, and counted for downstream bead based CD11b positive selection (StemCell Technologies, Vancouver, BC, Canada). The manufacturer’s protocol was followed for CD11b+ selection with two exceptions, 1) cells were placed on ice and mixed every five minutes for all incubations and 2) 10 mL of biotinylated antibody (BioLegend, San Diego, CA; 0.5 mg/ml, clone M1/70) and 100 mL of Selection Cocktail was added for every 5x10^7^ cells instead of per mL of sample. Purity of isolated Kupffer cells was determined by flow cytometry, as discussed below.

### Western blot

MDM, Kupffer cells, liver, thalamus, and parietal cortex were lysed with 1X RIPA buffer (Cell Signaling Technology, Danvers, MA) supplemented with 1X protease/phosphatase inhibitor (Cell Signaling Technology). Total protein concentrations were determined by Bradford Assay using the Bio-Rad Protein Assay Dye Reagent (Bio-Rad, Hercules, CA) following manufacturer instructions. Forty μg of total protein per sample was electrophoresed on a 4– 15% polyacrylamide gradient gel (Bio-Rad) and transferred to nitrocellulose membranes (Amersham Biosciences; Woburn, MA). Following membrane blocking in 3% BSA (Fisher Scientific) and 5% non-fat dried milk (Lab Scientific bioKEMIX, Danvers, MA), membranes were probed with primary antibodies specific to CYP3A4 (Thermo Fisher Scientific), PXR (Thermo Fisher Scientific), CAR (R&D Systems), GAPDH (Cell Signaling Technology), or SIGMAR1 (Cell Signaling Technology) overnight at 4°C. Membranes were probed with their appropriate HRP-conjugated secondary anti-rabbit (ABCAM, Cambridge, UK) or anti-mouse (ABCAM) antibody at room temperature for 1 hour. Protein expression was visualized via chemiluminescence (Revvity Health Sciences, Waltham, MA)) and quantified according to our established protocols^20–22^.

As a loading control, membranes were stripped with Restore Plus Western Blot Stripping Buffer (Thermo Fisher Scientific) and re-probed with antibody against β-actin HRP (Cell Signaling Technology, 8H10D10) for 1 hour at room temperature. Densitometric analysis was performed using ImageJ (version 1.53t, NIH, Bethesda, MD) to quantify band densities (pixels) for target proteins. Relative protein expression was calculated by normalizing each band to β-actin, with the vehicle-treated group set to 1, and fold changes determined relative to vehicle.

Due to the highly distinct biological profiles of Kupffer cells and whole liver tissue homogenates, that are primarily comprised of hepatocytes, traditional housekeeping proteins like GAPDH and β-actin exhibit significant cell-type-specific variability^23–26^. Thus, to ensure quantitative accuracy and mitigate the limitations of differing intracellular concentrations of traditional housekeeping protein expression, total protein loading was assessed. To accomplish this, membranes were incubated in Ponceau S solution (Sigma-Aldrich) for 30 seconds, briefly rinsed with distilled water to visualize total protein, and imaged for quantification. The stain was then completely removed following washed with dH_2_0 prior to blocking.

### Immunofluorescent microscopy

Immunostaining was performed using previously established protocols^20,27^. Primary human macrophages were fixed with 4% paraformaldehyde (Electron Microscopy Sciences, Hatfield, PA) for 15 minutes at room temperature and permeabilized with 0.01% Triton X-100 (Sigma-Aldrich, Burlington, MA). To visualize cell morphology, cells were stained with wheat germ agglutinin conjugated to Texas red (Fisher Scientific). To reduce nonspecific binding, cells were incubated with blocking buffer containing 5 mM EDTA (Sigma-Aldrich), 1% fish gelatin (Sigma-Aldrich), 1% immunoglobulin- free bovine serum albumin (Sigma-Aldrich), 1% heat-inactivated human serum AB (GeminiBio, Sacramento, CA) and 1% goat serum (Vector Laboratories, Newark, CA) in 1X DPBS (Gibco). Cells were probed overnight at 4 °C with a primary antibody specific to CYP3A4 (Thermo Fisher Scientific), followed by incubation with the appropriate Alexa Fluor 488-conjugated secondary antibody (Thermo Fisher Scientific) for one hour at room temperature. Following final washes, cells were mounted with Ibidi mounting medium containing DAPI as a counterstain to identify nuclei (Ibidi USA, Madison, WI). Fluorescence imaging was performed using the ECHO Revolution microscope (Echo, San Diego, CA) to evaluate CYP3A4.

### CYP3A4 Enzymatic Activity Determination

To evaluate the metabolic capacity of CYP3A4 in macrophages, PBMC were seeded at a density of 30,000-50,000 cells per well and differentiated to MDM as described above in black walled, clear bottom 96-well plates (Corning, Corning, NY). MDM were treated for 24 hours at 37°C in 5% CO₂ with cocaine hydrochloride (10 μM, NIDA Drug Supply Program), rifampicin (1 μM, Millipore Sigma), dexamethasone (10 μM, Millipore Sigma), norcocaine (10 μM, NIDA Drug Supply Program), benzoylecgonine (10 μM, NIDA Drug Supply Program), or dopamine (10 μM, Millipore Sigma). Treatment with vehicle was used as a control.

Following treatment, cells were washed twice with 1X DPBS (Gibco) and incubated with the CYP3A4 probe substrate BFC (2 μM; Sigma-Aldrich). BFC stock was prepared according to the manufacturer’s instructions in acetonitrile (0.0375%, Millipore Sigma) and diluted in phenol-red free macrophage media. To enhance CYP3A4-mediated metabolite formation and improve assay sensitivity, an NADPH-generating system was added and prepared fresh prior to each assay as described by Cheng et al^28^. The system consisted of 50 parts 10 mM NADP⁺ (Millipore Sigma), 50 parts 100 mM glucose-6-phosphate (Millipore Sigma), and 1 part 1 mg/mL yeast glucose-6-phosphate dehydrogenase (Millipore Sigma) containing 1.0 mM EDTA (Quality Biological Inc, Gaithersburg, MD) and 20% glycerol (v/v) (Millipore Sigma). During an incubation period of 90-120 minutes at 37°C, formation of the fluorescent metabolite HFC was quantified using a SpectraMax microplate reader (Molecular Devices, San Jose, CA) (excitation/emission: 405/535 nm). CYP3A4 velocity was determined as previously described^20^.

### Flow cytometry

To confirm purity of Kupffer Cell isolation, 1*10^6^ liver cells were obtained before and after CD11b positive selection, stained with antibodies for CD3 (SP34-2, Becton Dickinson [BD], Franklin Lakes, NJ) and CD11b (ICRF44, BioLegend, San Diego, CA) for 30 minutes on ice in FACS buffer (1% FBS (VWR, Radnor, PA) in DPBS (Gibco)), filtered with cell strainer caps (BD), and acquired on the BD FACS Lyric flow cytometer. At least 50,000 events were collected. Unstained, isotype matched antibodies, and fluorescence minus one staining were used as controls and to determine positivity for each antigen. Data were acquired using BD FACSuite software and analyzed using FlowJo v11 (FlowJo; Ashland, Oregon).

### Bulk RNA sequencing

MDM from six independent donors were treated with cocaine hydrochloride (10 μM, NIDA Drug Supply Program) or vehicle for 24 hours, cells lysed, and RNA isolation performed as previously described^22,29^. RNA was sent to the Emory National Biomedical Research Center Genomics Core to confirm purity and for library preparation, sequencing, and data analysis.

### CYP3A4 genotyping assay for single nucleotide polymorphisms (SNPs)

Genomic DNA was extracted and purified using QIAmp DNA blood mini kit (Qiagen, Germantown, MD). To evaluate CYP3A4 SNPs using TaqMan SNP genotyping assays (Fisher Scientific; *22 C__59013445_10, *18 C__27859823_20, *1G C__26201900_30, and *2 C__30634204_10) reactions were run in duplicate using TaqMan Genotyping Master Mix (Thermo Fisher Scientific) and nuclease free water (New England Biolabs, Ipswich, MA) according to the manufacturer’s protocol, and amplification was performed on the CFX Real-Time PCR System (Bio-Rad, Hercules, CA). Endpoint fluorescence data were used for allelic discrimination, and genotype calls were assigned using instrument software.

### LEGENDplex cytokine analysis

MDM were treated with cocaine hydrochloride (10 mM) or vehicle and after 24 hours supernatant was collected, aliquoted, and stored at -80°C. There were no freeze/thaw cycles before analysis. The LEGENDPlex human inflammation panel (BioLegend) was used to evaluate IL-1β, IFN-α2, IFN-γ, TNF-α, MCP-1, IL-6, CXCL8 (IL-8), IL-10, IL-12p70, IL-17A, IL-18, IL-23, and IL-33 according to manufacturer’s protocol and as previously described^20,22,30^. Cytokine and Chemokine concentrations were determined using a BD FACSLyric and analyzed using Qognit software version 2025-05-01.

### High-Content Imaging

MDM were fixed with 4% paraformaldehyde (Thermo Fisher Scientificfor 10 minutes at room temperature, permeabilized, incubated with blocking buffer (1% BSA (Fisher Scientific), 0.1% Tween 20 (Fisher Scientific), and 22.52 mg/mL glycine (Fisher Scientific) in 1X PBS) for 30 minutes. Cells were stained with anti-p24-Gag primary antibody (ATCC, Manassas, Virginia, USA), counterstained with DAPI (Thermo Fisher Scientific) and HCS CellMask Deep Red (Thermo Fisher Scientific), and high content imaging performed to evaluate HIV-infected p24+ cells as previously described^22^.

### HIV p24 Quantification

HIV p24 capsid antigen in MDM supernatants was quantified with an Alliance HIV-1 ELISA Kit (Perkin Elmer NEK050B001KT; Shelton, Connecticut, USA) according to the manufacturer’s protocol with a limit of detection of 6.25 pg/mL.

### Rhesus macaque experimental design

Animals and procedures in this study were approved by and in compliance with Emory University and Johns Hopkins Animal Care and Use Committees. Animal handling and euthanasia were conducted as stated under the NIH Guide for the Care and Use of Laboratory Animals and the USDA Animal Welfare Regulation. Animals were included in the study if they were healthy, adult rhesus macaques (*Macaca mulatta*) that were experimentally naïve and negative for the immunoprotective MHC class I alleles *Mamu-A*01*, *Mamu-B*08*, and *Mamu-B*13*. Macaques were pair-housed in the same facility to minimize any immunologic stress caused by being single-housed and given *ad libitum* access to standard monkey chow (Teklad Diet 2018; Indianapolis, Indiana, USA) and water. Rhesus macaque cohorts included at least n=2 per biological sex, n=4-5 total in this study, for each cohort to detect a 55% change in a dataset that demonstrates a standard deviation of 0.33 (power of 80%) and significance level of 0.05 (R statistical software). The first cohort included animals (N=4) who received a subcutaneous dose of 2.5 mg/kg dolutegravir (DTG, Viiv, London, England, UK), 20 mg/kg tenofovir disoproxil fumarate (TDF, Gilead, Foster City, California, USA), and 40 mg/kg emtricitabine (FTC, Gilead, Foster City, California, USA) daily for one month. The second cohort was comprised of macaques (N=5) who received cocaine hydrochloride at 0.5 mg/kg/day intravenously, 3 times weekly for one month followed by coadministration of the same antiretroviral regimen as cohort one for an additional month. The dose of cocaine was selected as a low-dose reinforcing concentration in self-administration models^31–39^. Animal weights were ∼5 kg and maintained throughout the study for all animals. For euthanasia, animals were sedated using ketamine and administered an overdose of sodium pentobarbital, according to the American Veterinary Medical Association guidelines. Upon euthanasia at ∼8 AM, tissue harvest was performed in the same manner for all animals and completed by 12 PM. Briefly, CSF and peripheral and chest blood were collected, followed by perfusion using 1X phosphate buffered saline. Tissues were subsequently collected, including brain, kidney, and liver. Brain was further dissected by region, including thalamus and parietal cortex.

### ART Determination

Forty-eighty mg of tissue were lysed in 70% LC-MS-grade methanol. FTC, TFV, FTC-TP, and TFV-DP were quantified in collaboration with the Clinical Pharmacology Analytic Laboratory at The Johns Hopkins University School of Medicine as previously described^21^.

### Statistical analysis

All statistical analyses were performed on data from four independent animals per macaque cohort or at least six independent donors for MDM experiments. Details regarding the number of independent experiments performed are included in all figure legends. Statistical analyses were performed using Prism software version 11 (GraphPad Software, Inc.; San Diego, California, USA). Statistical significance was determined by first performing a normality test to determine whether the data were normally distributed. If so, an ANOVA test was performed for unpaired multiple comparisons, and a two-tailed Student’s t-test for pairwise comparisons. If the data were not normally distributed, a Kruskal-Wallis test was performed for unpaired multiple comparisons, Friedman test was performed for paired multiple comparisons, Mann-Whitney was performed for unpaired pairwise data, and Wilcoxon tests for paired pairwise data. As the statistical test used in the respective panels often differed, the statistical analysis approach for each figure is detailed in the respective Figure Legends. *p<0.05, **p<0.01, ***p<0.001, and ****p<0.0001.

## Results

### Human Macrophages Possess Functional and Inducible CYP3A4 with High Levels of Inter-Individual Variability

To characterize macrophages as a site of extrahepatic drug metabolism, we first assessed whether they expressed the primary phase I drug metabolizing enzyme, CYP3A4. We performed Western blot to evaluate total protein abundance in MDM cell lysates from 20 donors. Representative blots revealed substantial inter-individual variability in basal CYP3A4 expression (**Figure 1A**). Densitometric analysis of CYP3A4 normalized to β-actin was performed, confirming constitutive enzyme expression with high variability among donors (**Figure 1B**). To confirm these observations, we performed immunofluorescent microscopy to visualize CYP3A4 in MDM from six donors. We identified ubiquitous cytoplasmic distribution and high inter-individual variability in the expression of CYP3A4 among all donors (**Figure 1C**).

**Figure 1:**
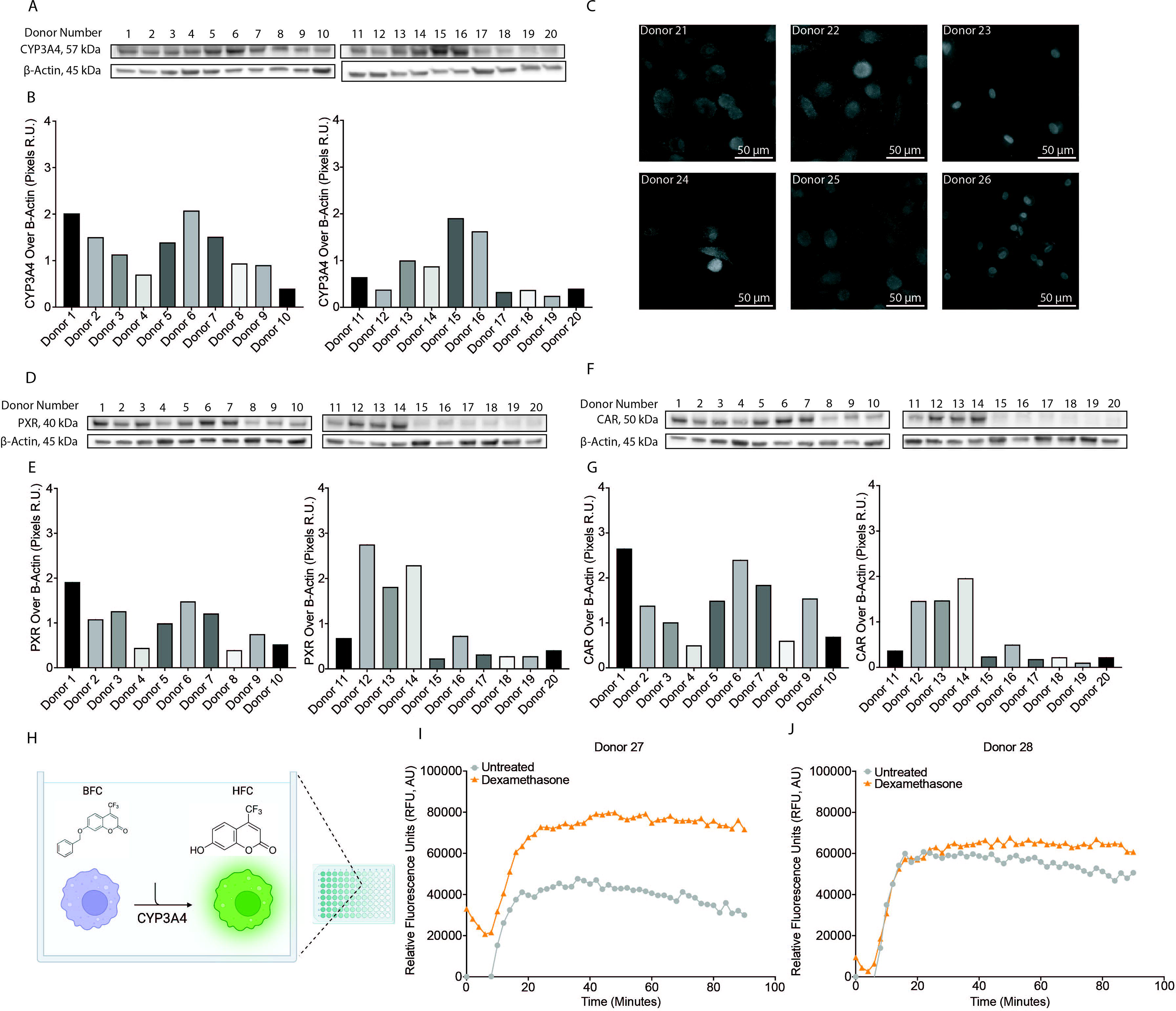
Human macrophages represent an extrahepatic compartment of CYP3A4-mediated drug metabolism. MDM were obtained from 20 independent donors and Western blot and densitometric analysis performed to evaluate (A-B) CYP3A4, (D-E) PXR and (F-G) CAR normalized to β-actin. (C) CYP3A4 was evaluated by immunofluorescent microscopy in MDM obtained from six independent donors. (H) Schematic representation of the microplate assay used to evaluate CYP3A4’s functional ability to catalyze the O-debenzylation of 7-benzyloxy-4-trifluoromethylcoumarin (BFC) to produce the fluorescent metabolite 7-hydroxy-4-trifluoromethylcoumarin (HFC). (I-J) CYP3A4’s catalytic activity was evaluated at baseline in untreated MDM (grey) or after dexamethasone (orange) treatment for 90 minutes and the relative fluorescence units (RFU) quantified in cells from two independent donors.

To probe the capacity for macrophages to undergo canonical CYP3A4 regulation, we next examined its well-established upstream regulators, PXR and CAR, that we hypothesized may be driving donor-dependent variations in CYP3A4. These ligand-activated nuclear receptors regulate the transcription of drug metabolizing enzymes, including CYP3A4, in response to xenobiotic exposure^40–42^. To address this, we performed Western blot to evaluate protein abundance in MDM lysates from the same 20 donors as in **Figure 1A**. Representative blots confirmed MDM PXR and CAR expression (**Figure 1D, 1F**) with densitometric analysis identifying high donor-specific variability for both proteins (**Figure 1E, 1G**). These findings demonstrate that MDM possess the regulatory machinery associated with CYP3A4-mediated drug metabolism, validating the importance of more comprehensively understanding their drug metabolic capacity.

To further characterize MDM as a functional site for extrahepatic drug metabolism, we measured basal metabolic activity using the fluorogenic CYP3A substrate, BFC, which is metabolized to the fluorescent product, HFC (**Figure 1H**). The rate of HFC production, indicated by fluorescent signal (relative fluorescence units, RFU), was measured over 90 minutes. Cells were pre-treated for 24 hours with vehicle or dexamethasone (10 μM), a potent CYP3A4 inducer. All donors had functional CYP3A4, but kinetic analysis revealed that the timing and enzymatic capacity of CYP3A4 to convert BFC to HFC was donor dependent. Donors with a robust response showed a higher baseline signal that increased rapidly within the first 20 minutes, where the fluorescent signal remained substantially higher than vehicle control for the entirety of the experiment (**Figure 1I**). In contrast, in donors with modest induction, the signal increased rapidly within the first 20 minutes, resulting in only a slight increase in fluorescence compared to vehicle (**Figure 1J**). The magnitude of basal CYP3A4 expression was not predictive of robust or more modest enzymatic induction by dexamethasone (data not shown). Collectively, these findings demonstrate that MDM possess functional and inducible CYP3A4 and may serve as a site of extrahepatic drug metabolism, possessing donor-dependent enzymatic activity.

### Rhesus Macaque Kupffer Cells Express CYP3A4

To determine whether our MDM findings extended to physiologic systems, we assessed CYP3A4 in rhesus macaque Kupffer cells. Liver was collected, enzymatic digestion performed, and Kupffer cells obtained by immunomagnetic isolation using antibodies with specificity for the pan-myeloid marker CD11b (**Figure 2A**). CD11b+ cells represented ∼90% of the total cells following CD11b selection, as compared to the ∼35% present before selection (**Figure 2B**). The remaining 10% of cells present following CD11b selection were CD3+ T cells. Now with a CD11b+ population enriched for Kupffer cells, we performed Western blot to evaluate CYP3A4 in comparison to whole liver homogenates. CYP3A4 was detectable in CD11b^+^CD3^-^ Kupffer cells from four independent rhesus macaques (**Figure 2C**), demonstrating the potential for macrophages to also have P450 expression *in vivo*. Interestingly, Kupffer cells also had relatively high expression of PXR, relative to the whole liver homogenate, demonstrating that they possess the canonical transcriptional machinery necessary to regulate CYP3A4 (**Figure 2C**). Of note, whole liver lysate is primarily comprised of hepatocytes that have substantially different ploidy, cellular metabolism, and size than Kupffer cells that make normalization difficult using traditional housekeeping proteins, including GAPDH and β-actin^23–25^. To this end, rather than perform quantitative analysis, we performed a total protein stain for illustrative purposes (**Figure 2D**)^26^. Together, these results demonstrate that liver macrophages express CYP3A4 and PXR, and are consistent with our MDM findings. Moreover, these data support further investigation into the contribution of macrophage populations, including those in the liver, to local drug metabolism.

**Figure 2:**
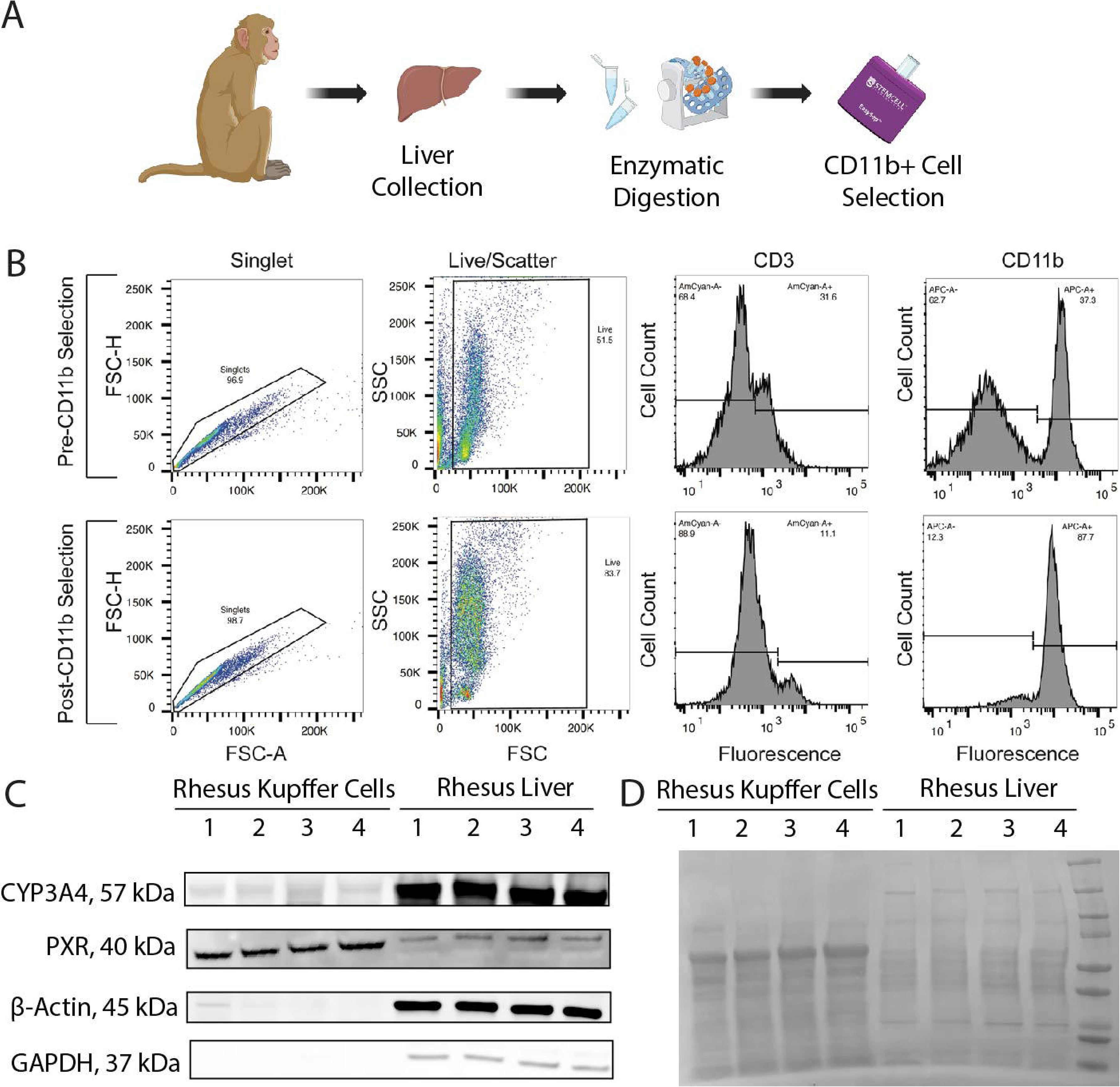
Rhesus Macaque Kupffer Cells Express CYP3A4. (A) Schematic of CD11b positive isolation from rhesus macaque liver. (B) Representative gating strategy used to evaluate cells before (top) and after (bottom) CD11b+ cell selection. Cells were first gated to evaluate singlets, followed by forward and side scatter characteristics, and ultimately evaluated for CD3 and CD11b positivity. (C) Western blot analysis of CYP3A4, PXR, β-actin, and GAPDH from lysates obtained from CD11b+ Kupffer cells and liver homogenate from four rhesus macaques. (D) Western blot membrane was stained with Ponceau S solution to visualize total protein content. Ladder used to identify molecular weight was loaded in the last well, and can be seen on the far right of the gel.

### Cocaine is a CYP3A4 Inducer

CYP3A4 enzymatic activity is readily modulated by a wide variety of xenobiotics that either increase or decrease its functional capacity, termed inducers and inhibitors, respectively. When this affects liver CYP3A4, inducers and inhibitors can perpetuate adverse drug:drug interactions that profoundly impact therapeutic pharmacologic profiles. We previously demonstrated that cocaine regulated CYP3A4 by PXR-dependent mechanisms at the human blood brain barrier (BBB)^20^. Further, cocaine has been established as a CYP3A4 substrate in human liver microsomes^43,44^. To evaluate whether macrophage CYP3A4 was also permissive to modulation by cocaine that may contribute to adverse drug:drug interactions, we characterized the impact of 24-hour treatment with cocaine, or vehicle in MDM obtained from 75 independent donors. Following treatment, Western blot was performed to evaluate CYP3A4 relative to β-actin. Unexpectedly, cocaine had a heterogenous effect where MDM from some individuals exhibited an increase in CYP3A4 (1.8 fold relative to vehicle, p<0.0001, Wilcoxon test), whom we termed “responders”, while others had a decrease or no change (0.9 fold relative to vehicle, p=0.0029, Wilcoxon test) that we classified as “non-responders” (**Figure 3A**). This dichotomous CYP3A4 expression “responsiveness” profile response was confirmed following densitometric analysis, where the fold change relative to vehicle control was determined, revealing the two distinct subgroups (**Figure 3B**). Interestingly, both subgroups were present in near equal numbers among the 75 donors as they each comprised ∼50% of our donor population (**Figure 3C-3D**).

**Figure 3:**
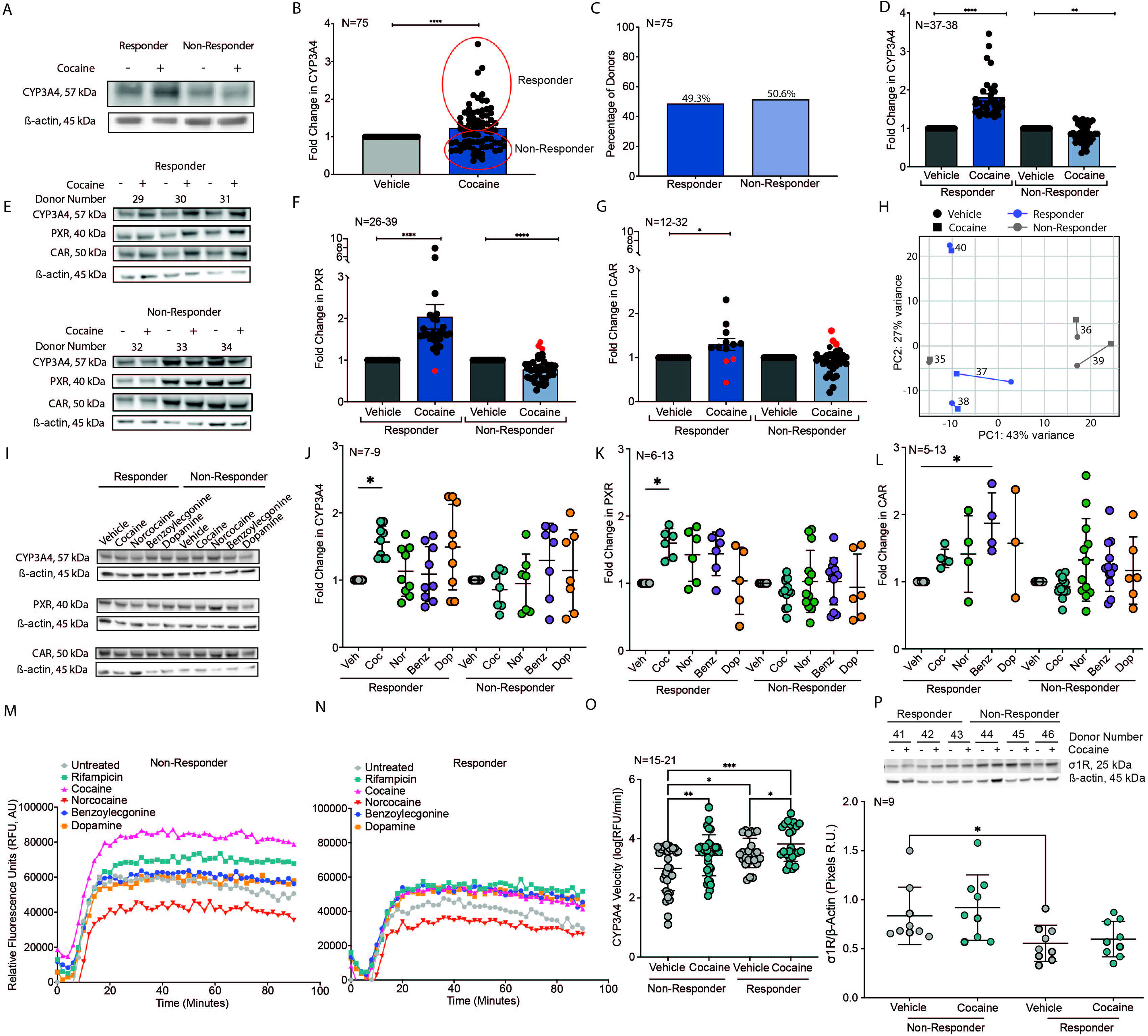
Cocaine Elicits Donor-Dependent CYP3A4 Expression Responses but Uniformly Enhances CYP3A4 Activity in Human Macrophages. MDM were treated with cocaine or vehicle for 24 hours and Western blot performed to evaluate CYP3A4 and β-actin. (A) Representative Western blot demonstrating the dichotomous CYP3A4 expression “responsiveness” profile in MDM following cocaine treatment. (B) MDM from 75 donors were obtained, treated with cocaine for 24 hours, Western blot performed to evaluate CYP3A4 and β-actin, and densitometric analysis performed to quantify the normalized fold change in CYP3A4/β-actin from cocaine (blue) treated cells, where vehicle (grey) was set to 1. Red circles identify the “Responder” and “Non-Responder” expression “responsiveness” profile based on the ability of cocaine to increase or decrease/cause no change in CYP3A4. Data are represented as mean ± SD. ****p<0.0001. Wilcoxon test. (C) The percentage of donors who had a “Responder” (dark blue) and “Non-Responder” (light blue) CYP3A4 expression profile following 24 hour cocaine treatment was determined. Data are represented as mean ± SD. (D) The normalized fold change in CYP3A4/β-actin from cocaine (blue) treated MDM, where vehicle (grey) was set to 1, for donors who had the “Responder” (dark blue) or “Non-Responder” (light blue) CYP3A4 expression profile. Data are represented as mean ± SD. **p<0.01. ****p<0.0001. ANOVA. (E) Representative Western blot evaluating CYP3A4, PXR, CAR, and β-actin for six donors who had a “Responder” or “Non-Responder” CYP3A4 expression profile following 24 hour treatment with cocaine or vehicle. Densitometric analysis was performed and the normalized fold change in (F) PXR/β-actin or (G) CAR/β-actin from cocaine (blue) treated MDM, where vehicle (grey) was set to 1, for donors who had the “Responder” (dark blue) or “Non-Responder” (light blue) CYP3A4 expression profile. Red dots indicate donors who had discordant PXR and CAR fold change following cocaine treatment relative to CYP3A4. Data are represented as mean ± SD. *p<0.05. ****p<0.0001. ANOVA. (H) Principal Component Analysis (PCA) plot of RNA-seq expression profiles of RNA obtained from six-independent MDM donors treated cocaine (square) or vehicle (circle) who had the “Responder” (blue) or “Non-Responder” (grey) CYP3A4 expression profile. (I) Representative Western blot evaluating CYP3A4, PXR, CAR, and β-actin for donors who had a “Responder” or “Non-Responder” CYP3A4 expression profile following 24 hour treatment with cocaine, norcocaine, benzoylecgonine, dopamine, or vehicle. Densitometric analysis was performed and the normalized fold change in (J) CYP3A4/β-actin or (K) PXR/β-actin, or (L) CAR/β-actin from MDM following cocaine (Coc, teal), norcocaine (Nor, green), benzoylecgonine (Benz, purple) or dopamine (Dop, orange) where vehicle (grey) was set to 1, for donors who had the “Responder” or “Non-Responder” CYP3A4 expression profile. Data are represented as mean ± SD. *p<0.05. ANOVA. Relative fluorescence units (RFU) quantifying BFC conversion to HFC over 90 minutes for MDM from representative donors who had the (M) “Non-Responder” or (N) “Responder” CYP3A4 expression profile following 24 hour treatment with rifampicin (teal square), cocaine (pink triangle), norcocaine (red downward triangle), benzoylecgonine (blue circle), dopamine (orange square), or vehicle (grey circle). (O) Log-transformed CYP3A4 velocity following vehicle (grey) or cocaine (teal) treatment from donors who had the “Non-Responder” or “Responder” CYP3A4 expression profile. Data are represented as mean ± SD. *p<0.05. **p<0.01. ***p<0.001. ANOVA. (P) Representative Western blot evaluating σ1R and β-actin for 18 donors who had a “Responder” (N=9) or “Non-Responder” (N=9) CYP3A4 expression profile following 24 hour treatment with cocaine or vehicle. Densitometric analysis was performed and the normalized fold change in σ1R /β-actin from cocaine (teal) treated MDM, where vehicle (grey) was set to 1, for donors who had the “Responder” or “Non-Responder” CYP3A4 expression profile. Data are represented as mean ± SD. *p<0.05. ANOVA.

To elucidate whether PXR and CAR were reflective of the dichotomous CYP3A4 expression “responsiveness” profile, we next evaluated the impact of cocaine on CYP3A4 protein expression. Following 24 hours of cocaine or vehicle treatment, PXR and CAR were measured by Western blot, and densitometric analysis performed relative to β-actin (**Figure 3E**). In “responders”, cocaine treatment significantly increased PXR (1.9 fold relative to vehicle, p<0.0001, Wilcoxon test, **Figure 3F**) and CAR (1.3 fold relative to vehicle, p=0.0342, ANOVA test, **Figure 3G)**, whereas in “non-responders” it significantly decreased PXR (0.8 fold relative to vehicle, p<0.0001, ANOVA test, **Figure 3G**), while CAR showed no significant change (0.9 fold relative to vehicle, p=0.2583, Wilcoxon test, **Figure 3F**). Cocaine promoted the same effect for CYP3A4, PXR, and CAR for almost all donors, wherein all proteins were similarly increased in “responders” or showed no change or a decrease in “non-responders”. The minority of individuals (4/65 for PXR and 6/44 for CAR) for whom this did not occur are indicated in red (**Figure 3F-G**). Collectively, these results suggest that cocaine likely modulates macrophage CYP3A4 through canonical PXR and CAR pathways; however, the variability in donor response suggests that additional regulatory factors may also be involved.

To further characterize the regulatory drivers behind the dichotomous effect of cocaine on the MDM CYP3A4 expression “responsiveness” profile, we performed bulk RNA sequencing (RNA-seq). Following treatment with cocaine or vehicle for 24 hours, we found that MDM transcriptomic responses were variable and not associated with the “responsiveness” profile (**Figure 3H**). Cocaine promoted a large impact on the transcriptome for some individuals (donors 37 and 39) while, unexpectedly, caused very little changes for others (donors 35, 38, and 40). We found no association with cocaine responsiveness and basal CYP3A4 levels among donors by RNA-seq and qRT-PCR, as the transcript was lowly expressed, or Western blot (data not shown). Similarly, donor demographic factors were not predictive of cocaine responsiveness, including biological sex (**Figure 4**), cytokine profiles in MDM supernatant following cocaine treatment, and CYP3A4*22, CYP3A4*18, or CYP3A4*1G genotype^45^ (data not shown).

**Figure 4:**
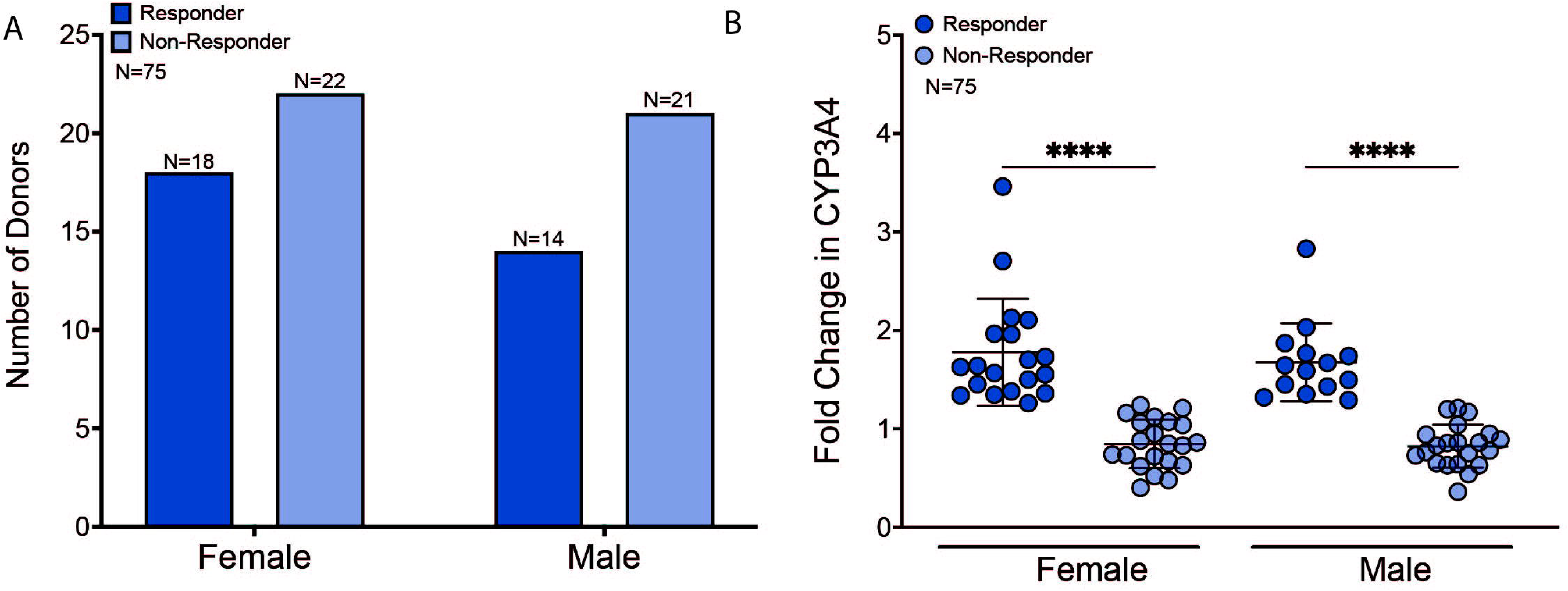
Biological Sex is not Associated with the Dichotomous CYP3A4 Expression Profile. Data were disaggregated by biological sex for 75 independent MDM donors according to their “Responder” (dark blue) or “Non-Responder” (light blue) CYPA4 expression profile following 24 hour cocaine or vehicle treatment. (A) The number of female and male donors associated with each CYP3A4 expression profile was enumerated. (B) The normalized fold change in CYP3A4/β-actin from cocaine treated MDM, where vehicle (grey) was set to 1, was determined for female and male donors with the “Responder” (dark blue) or “Non-Responder” (light blue) CYPA4 expression profile. Data are represented as mean ± SD. ****p<0.0001. ANOVA.

As the transcriptome and donor demographic factors were not indicative of the dichotomous cocaine CYP3A4 expression “responsiveness” profile, we queried whether its effects on CYP3A4 were specific or if they also occurred by cocaine metabolites or dopamine – the neurotransmitter that mediates cocaine’s addictive and reinforcing effects. Macrophages were treated for 24 hours with cocaine, its minor metabolite norcocaine, its major metabolite benzoylecgonine, dopamine (all at 10 μM), or vehicle. CYP3A4, PXR, and CAR were evaluated by Western blot and normalized to β-actin (**Figure 3I**). We evaluated CYP3A4 in MDM from 16 donors, classifying 9 individuals as “responders” and 7 as “non-responders” **(Figure 3J**). We evaluated PXR in MDM obtained from 19 donors, classifying 6 individuals as “responders” and 13 as “non-responders” (**Figure 3K**). Finally, we characterized CAR using cells from 18 MDM donors, classifying 5 individuals as “responders” and 13 as “non-responders” (**Figure 3L**). In “responders”, cocaine significantly increased CYP3A4 (1.6 fold relative to vehicle, p=0.03, ANOVA; **Figure 3J**) and PXR (1.6 fold relative to vehicle, p=0.02 ANOVA; **Figure 3K**), whereas CAR had an increase that was not statistically significant due to a lower magnitude and degree of freedom (1.4 fold relative to vehicle, p=0.6, ANOVA; **Figure 3L**). In contrast, there was no consistent trend for any of the cocaine metabolites or dopamine. Together, these findings indicate that the ability to regulate CYP3A4, PXR, and CAR are specific to cocaine, rather than its metabolites or dopamine, motivating further evaluation of CYP3A4 functional activity under these conditions.

Surprised by the unexpected dichotomous CYP3A4 expression “responsiveness” profile, we next evaluated cocaine’s impact on CYP3A4 catalytic activity. MDM from 36 donors were incubated with the fluorogenic CYP3A substrate BFC, and HFC production was measured as RFU over 90 minutes. Cells were pre-treated for 24 hours with cocaine, benzoylecgonine, norcocaine, dopamine (all at 10 μM), a strong CYP3A4 inducer rifampicin (2 μM) as a positive control, or vehicle. To our surprise, cocaine induced a similar effect on CYP3A4 catalytic activity for all donors – irrespective of their CYP3A4 expression profile (**Figure 3M-N**). Indeed, MDM from all donors had functional, basal CYP3A4 activity following vehicle treatment, which was increased with rifampicin. Interestingly, cocaine increased CYP3A4 activity for both “responders” and “non-responders” to a comparable or even higher extent than rifampicin (**Figure 3M-N**). Norcocaine, the CYP3A4-mediated metabolite of cocaine, is a known CYP3A4 inhibitor that self-limits further cocaine metabolism through a negative feedback loop^43,44,46^. Accordingly, and as expected, norcocaine decreased CYP3A4 activity for both “responders” and “non-responders”. Benzoylecgonine and dopamine had minimal and inconsistent effects where they caused no change in CYP3A4 activity for some donors (**Figure 3M**) while increasing activity for others (**Figure 3N**). However, this was unrelated to cocaine responsiveness as benzoylecgonine and dopamine had similar effects on CYP3A4 activity for both “responders” and “non-responders” (data not shown). Kinetic analysis was performed to evaluate CYP3A4 velocity, or the rate at which HFC-associated fluorescence occurred (RFU/min), according to our established method^20^. Cocaine significantly increased CYP3A4 velocity (RFU/min) for both “responders” (cocaine: 3.8±0.6 RFU/min vs. vehicle: 3.5±0.5 RFU/min, p=0.0118, one-way ANOVA) and “non-responders” (cocaine: 3.4±0.7 RFU/min vs. vehicle: 2.9±0.8 RFU/min, p=0.001, ANOVA), compared to vehicle (**Figure 3O**). However, the magnitude of the effect was significantly larger for “non-responders” as they had slower, basal CYP3A4 velocity compared to “responders” (p=0.0143, ANOVA).

In our final efforts to identify the mechanism underlying the dichotomous CYP3A4 expression “responsiveness” profile, we evaluated the sigma-1 receptor (σ1R), a polyfunctional intracellular chaperone for whom cocaine is an agonist and mediates its reinforcing behaviors^45,47–64^. We evaluated the impact of cocaine on σ1R in MDM from 18 donors, classifying 9 individuals as “responders” and 9 as “non-responders” by performing Western blot and densitometric analysis following 24 hours of treatment (**Figure 3P**). “Responders” had significantly lower basal levels of σ1R as compared to “non-responders” (0.55±0.18 pixels as compared to 0.84±0.29 pixels, p=0.03, ANOVA), which were maintained following cocaine treatment. These findings suggest an inverse relationship between σ1R and CYP3A4 expression “responsiveness” profile wherein decreased basal σ1R impacts cocaine-σ1R signaling in “responders”, that facilitates increased CYP3A4 expression and increased enzymatic activity, as compared to “non-responders”. Collectively, these findings demonstrate that cocaine is a CYP3A4 inducer in macrophages that may mediate its donor-dependent effects through σ1R.

### Cocaine Induces Variable and Donor-Dependent Changes in Genes Associated with Xenobiotic-Mediated Transport and Metabolic Responses in Human Macrophages

As bulk RNA-seq on cocaine- or vehicle-treated MDM showed large variability in transcriptome responses that were independent of the CYP3A4 expression “responsiveness” profile, we further elucidated donor-specific responses to cocaine by evaluating genes canonically associated with drug transport, phase 1 and phase 2 metabolism, and xenobiotic response regulation from the RNA-seq datasets. Genes of interest were arranged in heatmaps to compare expression between cocaine- and vehicle-treated MDM across “responders” and “non-responders" (**Figure 5**). Importantly, normalized counts for each gene were used instead of fold changes controlled to vehicle, adjusted p-values, or z-scores to highlight individual gene expression per donor and treatment. Consistent with our prior analysis, cocaine-induced changes in gene expression varied across donors regardless of responder classification. The genes with the highest expression corresponded to drug transporters involved in influx: OATP2B1 (*SLCO2B1*) and ENT1 (*SLC29A1*), and efflux: MRP1 (*ABCC1*) and MRP3 (*ABCC3*) (**Figure 5A**); while those with medium expression included the intracellular cocaine targets retinoid X receptor alpha (*RXRA*) and σ1R (*SIGMAR1*), as well as the efflux transporters BCRP (*ABCG2*) and MRP4 (*ABCC4*) (**Figure 5B**). Interestingly, the phase 1 metabolizing enzyme CYP3A5 and our transcription factors of interest, PXR (*NR1I2*) and CAR (*NR1I3*), had low expression across all donors, along with the efflux transporters P-gp (*ABCB1*) and MRP2 (*ABCC2*) (**Figure 5C**). Additional selected genes associated with drug metabolism and transport, xenobiotic response regulation, and dopamine-monoamine pathways also exhibited variable donor-dependent responses to cocaine (**Supplemental Figure 1**). Surprisingly, CYP3A4 expression was below the level of detection for all donors and treatments, consistent with our qPCR analyses above (data not shown). Taken together, these findings support our observation that cocaine-induced changes in transcriptome do not determine cocaine “responsiveness” in human MDM.

**Figure 5:**
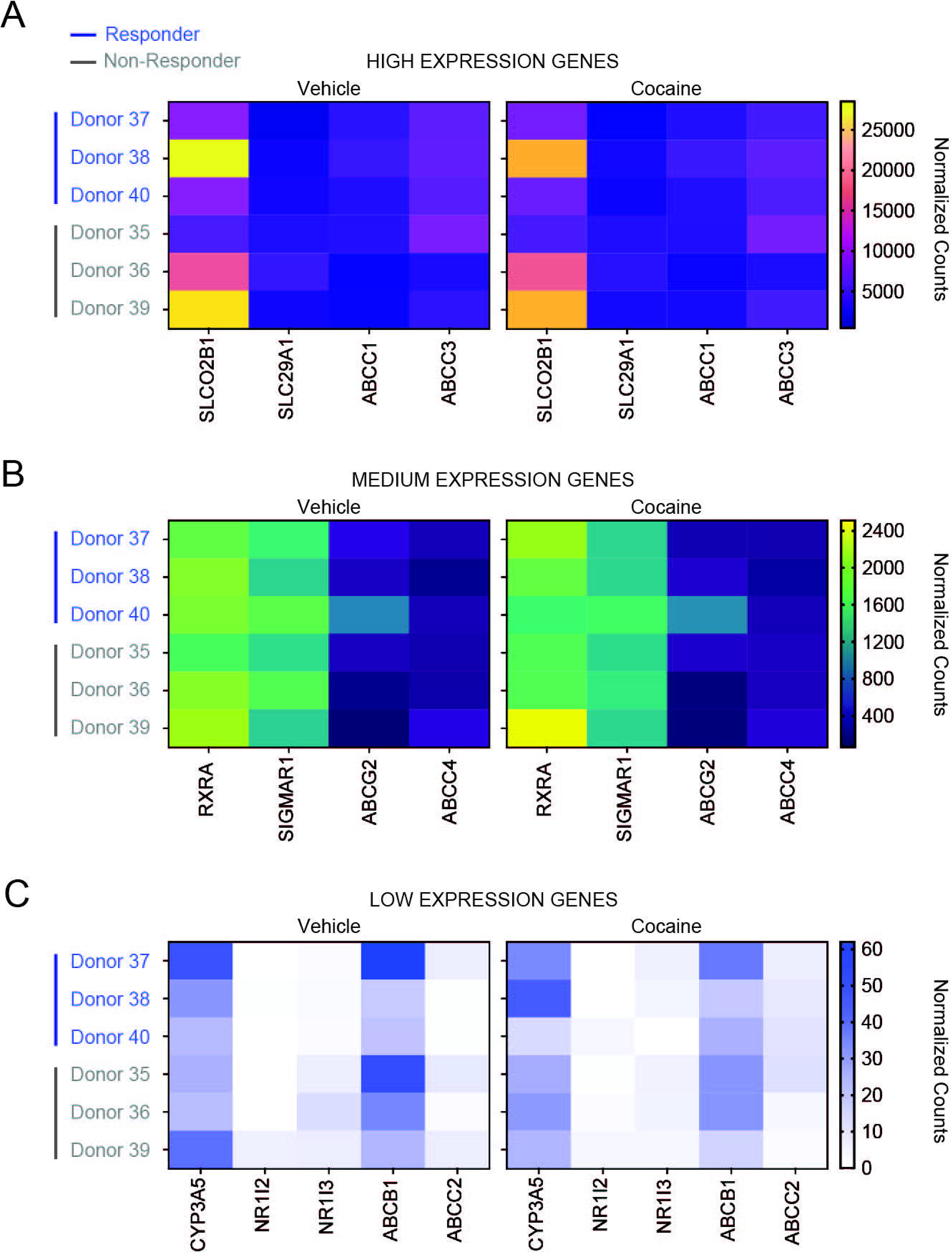
Drug transport and xenobiotic response gene expression varies across donors but does not distinguish CYP3A4 responders from non-responders following cocaine exposure. Heatmaps show normalized transcript counts for genes associated with drug transport and xenobiotic response across six donors under vehicle- and cocaine-treated conditions. Donors are ordered consistently across panels, where those who had the CYP3A4 “Responder” expression profile indicated in blue (Donors 37, 38, and 40) and donors with the “Non-responder” CYP3A4 expression profile indicated in gray (Donors 35, 36, and 39). Genes were separated according to expression abundance to facilitate visualization of differences across donors: (A) high-expression genes, *SLC6A11, SLC30A1, ABCC1, and ABCC3*; (B) medium-expression genes, *RXRA, SIGMAR1, ABCC2, and ABCC4*; and (C) low-expression genes, *CYP3A5, NR1I2, NR1I3, ABCB1, and ABCG2*. Color scales represent normalized transcript counts and are independently scaled for each expression group. Color intensity represents normalized counts according to the scale shown adjacent to each heatmap.

### Cocaine Participates in Adverse Drug:Drug Interactions That Increases CYP3A4 and PXR and Decreases Antiretroviral Therapy Tissue Distribution

Intrigued by the implications of our MDM findings, we next wanted to evaluate cocaine’s potential for adverse drug:drug interactions *in vivo*. We focused on two first-line antiretroviral therapies, tenofovir (TFV) and emtricitabine (FTC), used for HIV treatment and prevention as macrophages are an important cellular target for HIV known to have unique pharmacologic mechanisms that impact antiretroviral therapy intracellular concentrations^65,66^ .This study included two groups of rhesus macaques, evenly divided by sex. Group 1 (N=4) received an antiretroviral therapy regimen consisting of TFV and FTC for one month while Group 2 (N=5) received the same regimen in the presence of cocaine (**Figure 6A**). At the end of the study, liver, kidney, cerebellum, frontal cortex, and thalamus were obtained and concentrations of TFV and FTC along with their phosphorylated active metabolites, TFV-diphosphate (TFV-DP) and FTC-triphosphate (FTC-TP) were determined by LC-MS. While ART and their active metabolites were found throughout the body for Group 1 animals, concentrations were below the limit of quantitation (BLQ) for all animals in Group 2, indicating cocaine hindered tissue ART distribution and biotransformation (**Table 1**). We determined that this may have occurred, at least in part, due to upregulation of CYP3A4 and PXR as they were significantly increased in both thalamus (CYP3A4: 1.1±0.2 vs 0.8±0.1 pixels, p=0.0476, Mann-Whitney test; PXR: 1.0±0.2 vs 0.6±0.08 pixels, p=0.0159, Mann-Whitney test) and parietal cortex (CYP3A4: 1.2±0.2 vs 0.6±0.05 pixels, p=0.0159, Mann-Whitney test; PXR: 1.1±0.2 vs 0.7±0.03 pixels, p=0.0159, Mann-Whitney test) of Group 2 rhesus macaques that were administered ART with cocaine, compared to Group 1 animals that did not receive cocaine (**Figure 6B-D**). Although TFV and FTC are not CYP3A4 substrates, the ability of cocaine to increase CYP3A4 and PXR may have contributed to decreased ART concentrations through broader phase I and phase II mechanisms, including modulation of drug transporters consistent with our previous BBB findings^20^. These findings are of major importance as cocaine is already established to increase HIV replication^67^. Indeed, our MDM findings confirm this premise as we found that MDM infected with HIV_ADA_ in the presence of cocaine had increased supernatant concentrations of the capsid antigen, p24 (**Figure 6E-G**). Together, our findings identify macrophages as a relevant source of extrahepatic CYP3A4-mediated metabolism vulnerable to adverse drug:drug interactions that have the potential for substantial clinical implications.

**Figure 6:**
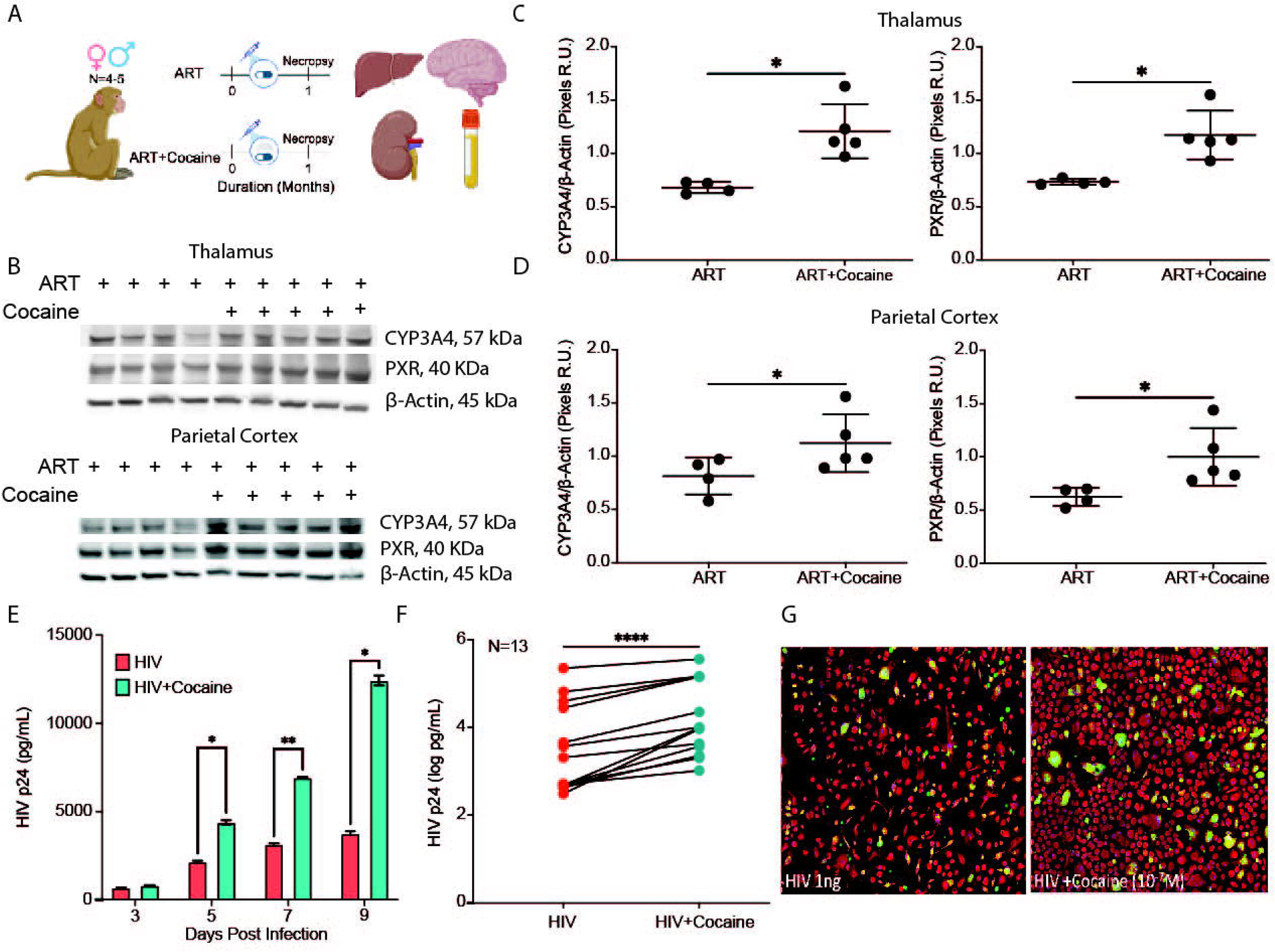
Cocaine alters ART drug disposition-associated pathways *in vivo* and enhances HIV replication in MDM. (A) Schematic of the rhesus macaque study design where animals received an ART regimen including TFV and FTC either alone (ART; N=4) or in combination with cocaine (ART + cocaine; N=5). Following one month of treatment, tissues were collected at necropsy for ART quantification by LC-MS and drug metabolism-associated proteins. (B) Representative Western blots of CYP3A4 and PXR in thalamus (top) and parietal cortex (bottom) from rhesus macaques receiving ART alone or ART+cocaine. β-Actin served as the loading control. (C, D) Densitometric analysis of CYP3A4 (C) and PXR (D) normalized to β-actin in thalamus (top) and parietal cortex (bottom). Data are presented as mean ± SD. *p < 0.05. Mann-Whitney test. (E) Representative HIV p24 determination in supernatants from HIV-1_ADA_-infected MDM cultured in the presence (teal) or absence (red) of cocaine 9 days post-infection. (F) HIV p24 concentrations in supernatants from HIV-1_ADA_-infected MDM from 13 independent donors in the presence (teal) or absence (red) of cocaine at the experimental endpoint, with matched samples connected by lines. ****p<0.0001. Wilcoxon test. (G) Representative high content images of HIV-1_ADA_-infected MDM cultured in the absence (left) or presence (right) of cocaine where green indicates the p24 capsid antigen. Cell morphology was determined using CellMask (red).

**Table 1.**
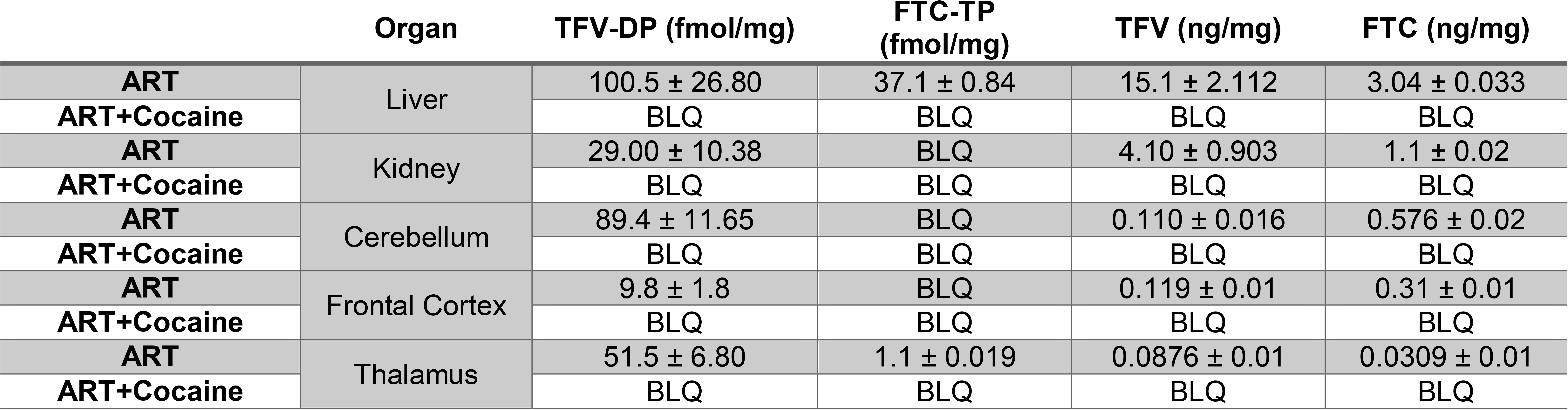
Cocaine Decreases ART and Active Metabolite Concentrations in Rhesus Macaque Tissues.

## Discussion

Our findings expand the conventional view of CYP3A4-mediated drug metabolism by identifying macrophages as a functionally active and highly heterogeneous extrahepatic metabolic compartment. Primary human macrophages exhibited measurable CYP3A4 activity with substantial interindividual variability, wherein CYP3A4 transcript, protein abundance, and catalytic function were poorly concordant. This distinction became particularly apparent following cocaine exposure: despite divergent donor-dependent effects on CYP3A4 protein levels, cocaine consistently increased catalytic activity across donors, to levels comparable to or exceeding rifampicin. The detection of CYP3A4 in rhesus macaque Kupffer cells further extended these observations beyond *in vitro* differentiated human macrophages. Moreover, cocaine-associated changes in ART tissue disposition that coincided with increased CYP3A4 and PXR levels *in vivo* demonstrated broader pharmacologic relevance. Together, these findings establish macrophages as an underrecognized component of extrahepatic drug metabolism, identify interindividual variation in CYP3A4 function that cannot be predicted from enzyme abundance alone, and provide evidence for substance use-mediated drug:drug interactions.

Drug metabolism is traditionally conceptualized as a function of the liver and intestine, where absorption, distribution, metabolism, and excretion processes influence systemic drug exposure and clearance. However, this paradigm does not fully capture metabolism occurring locally within distinct cellular niches. Macrophages represent a particularly important extrahepatic metabolic compartment because they reside in virtually every tissue and can express drug metabolizing enzymes capable of altering compounds in their intracellular microenvironment^68–71^. In this context, the significance of macrophage-mediated metabolism does not lie in its contribution to systemic clearance, but rather in its capacity to shape local cellular pharmacokinetics, including the concentration, persistence, and metabolic fate of drugs within macrophages themselves. This distinction is especially relevant for compounds whose therapeutic activity depends on achieving and maintaining sufficient intracellular exposure, including ART. Consequently, macrophage-specific metabolism could generate intracellular drug exposures that differ substantially from those predicted by plasma pharmacokinetics, *in silico* determinations, or hepatic metabolism alone. Defining macrophages as an important metabolic compartment therefore expands the conventional framework of drug disposition from systemic pharmacokinetics to cell-specific exposure, with direct implications for the design, dosing, and interpretation of macrophage-targeted pharmacotherapies whose efficacy depends on intracellular concentrations.

A striking finding of our study was the substantial interindividual variability observed across the 100 evaluated MDM donors. Heterogeneity in CYP3A4 was evident not only in basal expression, but also in the magnitude and direction of the response to cocaine, demonstrating a distinct expression-responder phenotype in which some individuals exhibited considerably greater CYP3A4 induction than others. Importantly, the mechanisms underlying this variability remain unclear. The tested SNPs did not account for the observed differences in expression responses, nor were these differences explained by sex, cytokines, cocaine metabolites, dopamine, or broader transcriptomic features evaluated in this study. Together, these findings suggest that MDM interindividual CYP3A4 variability may reflect biological determinants not readily captured by genetic, demographic, inflammatory, or transcriptional measures. Additional factors, including other measures of genetic variation, epigenetic regulation, environmental exposures, cellular activation state, and regulatory processes operating beyond steady-state transcript abundance, warrant investigation. Despite the uncertainty regarding how this donor-dependent variability occurred, our findings may have important implications for personalized medicine. If individuals differ substantially in the magnitude of their biological response to cocaine, or other CYP3A4 modulators, such variation could contribute to differences in disease susceptibility, progression, or therapeutic response. Collectively, the implications of our findings may ultimately inform approaches to patient stratification and individualized intervention.

We had the interesting finding that the effects of cocaine on CYP3A4 protein abundance did not parallel its effects on catalytic activity, identifying an important disconnect between CYP3A4 protein expression and function. Whereas cocaine altered CYP3A4 protein levels in a donor-dependent manner, CYP3A4 activity increased consistently across all evaluated donors. This divergence was particularly evident in individuals for whom cocaine caused a decrease or no increase in CYP3A4 expression: they exhibited relatively low basal CYP3A4 activity yet showed the largest functional induction following cocaine treatment. Thus, our data demonstrate that CYP3A4 protein abundance alone did not predict the magnitude of the response to cocaine. Additionally, these findings suggest that CYP3A4 protein expression-activity discordance may occur by post-transcriptional or post-translational regulatory mechanisms that influence enzyme stability, localization, catalytic efficiency, or interactions with regulatory proteins. Interestingly, our identification of an association between σ1R basal expression and CYP3A4 catalytic activity suggests that σ1R signaling may contribute to interindividual responses to cocaine-induced CYP3A4 regulation. Together, our findings caution against inferring CYP3A4 activity from protein expression alone and instead point to a functionally regulated response particularly pronounced in donors with low basal activity, especially in the context of local extrahepatic metabolism.

Similarly, our findings highlight an apparent disconnect between transcriptional and functional measures of macrophage CYP3A4 capacity. Unlike the gold-standard mRNA routinely used as a proxy for enzyme activity^72–75^, CYP3A4 mRNA remained below the limit of detection in both qPCR and transcriptomic analyses despite finding measurable CYP3A4 protein expression and enzymatic activity, demonstrating that transcript abundance does not necessarily predict the presence of a functional metabolic phenotype. This discordance may reflect low transcript abundance, high transcript turnover, differences in mRNA and protein stability, or post-transcriptional mechanisms that maintain protein levels despite limited detectable transcription. This premise has already been established in hepatocytes through miRNA regulatory processes^45^, alternative processing of the 3’-untranslated region (UTR) that leads to CYP3A4 transcripts with less mRNA stability^76^, post-transcriptional endoplasmic reticulum-associated E3 ubiquitin ligase regulation that controls CYP3A4 proteolysis and protein turnover, CYP3A4 phosphorylation^77^, and modulation by ubiquitin-proteasome and autophagy that regulate steady-state accumulation of CYP3A4 protein independently of transcription^45^. However, these mechanisms of CYP3A4 regulation are yet to be established in macrophages.

Our work may have important translational implications for HIV persistence and treatment in the setting of cocaine. Macrophages represent both a biologically relevant cellular compartment for HIV and a pharmacologic site in which adequate ART exposure is necessary to sustain antiviral activity ^78–84^. Notably, cocaine exposure in our rhesus macaque model was associated with markedly reduced tissue ART concentrations, suggesting that cocaine may alter the pharmacologic environment within tissues in ways that could compromise local drug exposure despite systemic therapy. Indeed, cocaine was associated with increased CYP3A4 and PXR expression, supporting broader cocaine-mediated remodeling of xenobiotic disposition pathways *in vivo*, including transport, renal handling, tissue uptake, intracellular phosphorylation/dephosphorylation, nucleotide/nucleoside metabolism, tissue distribution, or other pharmacokinetic processes. To this end, the observed changes in TFV and FTC tissue concentrations should not be attributed directly to CYP3A4 activity, as they are not metabolized by this P450. Rather, when considered together with the broader changes observed in xenobiotic metabolizing and disposition, these findings suggest that cocaine may have a more general impact on PXR-mediated transport, metabolism, distribution, and cellular disposition pathways. This pharmacologic effect may converge with cocaine-associated increases in macrophage HIV replication to create a potential “two-hit mechanism”, in which increased macrophage permissiveness to HIV is accompanied by reduced ART exposure at relevant tissue sites. Such convergence could be particularly consequential within pharmacologically protected compartments, including the brain, that represents an important HIV reservoir. This interpretation is consistent with our prior BBB findings demonstrating that cocaine perturbs PXR-mediated mechanisms governing drug disposition at the neurovascular interface^20^. Collectively, our present results extend that framework by suggesting that cocaine-associated disruption in ART distribution may also occur within tissue and cellular compartments relevant to HIV persistence.

Several considerations temper interpretation of our findings and highlight priorities for future investigation. Although BFC turnover supports CYP3A-associated catalytic activity, it does not establish CYP3A4 as the exclusive contributor to HFC formation. Additionally, blood-derived MDM may incompletely reflect the phenotypic and metabolic heterogeneity of tissue-resident macrophages. Likewise, although the liver macrophage preparation was ∼90% CD11b+, residual T-cell contamination may impact our findings. It should also be considered that our associations involving σ1R provide a basis for mechanistic investigation rather than evidence of causality. Moreover, RNA-seq from six donors limits resolution of molecular determinants underlying the heterogeneous phenotype. Similarly, analysis of our selected SNP variants had a similar limited capture of CYP3A4 genetic diversity. Finally, the independent macaque cohorts used in our study limit causal pharmacokinetic inference. Thus, the consistently BLQ tissue ART concentrations in cocaine-exposed animals warrant further investigation that, based on a single terminal time point, cannot distinguish among potential pharmacokinetic mechanisms.

Despite these considerations, our work challenges the liver-centric paradigm of drug metabolism by identifying macrophages as a functionally active, extrahepatic site of CYP3A4-mediated biotransformation. Using primary human macrophages and rhesus macaque tissues, we identify substantial interindividual variability and a critical disconnect between CYP3A4 expression and catalytic activity. Cocaine further highlights this complexity, uniformly enhancing CYP3A4 for all evaluated donors’ activity despite discordant mRNA and protein expression levels and coinciding with reduced ART tissue concentrations *in vivo*. These findings redefine macrophages as active determinants of tissue pharmacology and establish a framework for understanding how immune cell metabolism, substance use, and individual variability may converge to shape drug exposure, efficacy, and drug-drug interactions.

## Abbreviations

ABCG2: ATP-binding cassette subfamily G member 2
ART: antiretroviral therapy
BBB: blood-brain barrier
BCRP: breast cancer resistance protein
BFC: 7-benzyloxy-4-trifluoromethylcoumarin
BLQ: below the limit of quantitation
BSA: bovine serum albumin
CAR: constitutive androstane receptor
CNS: central nervous system
CYP: cytochrome P450
CYP3A4: cytochrome P450 3A4
DAPI: 4′,6-diamidino-2-phenylindole
DMEM: Dulbecco’s modified Eagle medium
DPBS: Dulbecco’s phosphate-buffered saline
DTG: dolutegravir
EDTA: ethylenediaminetetraacetic acid
ELISA: enzyme-linked immunosorbent assay
ENT1: equilibrative nucleoside transporter 1
FBS: fetal bovine serum
FTC: emtricitabine
FTC-TP: emtricitabine triphosphate
GAPDH: Glyceraldehyde 3-phosphate dehydrogenase
HFC: 7-hydroxy-4-trifluoromethylcoumarin
HIV: human immunodeficiency virus
LC-MS: liquid chromatography-mass spectrometry
M-CSF: macrophage colony-stimulating factor
MDM: monocyte-derived macrophages
MHC: major histocompatibility complex
MRP: multidrug resistance-associated protein
NADP⁺: nicotinamide adenine dinucleotide phosphate
NADPH: reduced nicotinamide adenine dinucleotide phosphate
OATP2B1: organic anion transporting polypeptide 2B1
PBMC: peripheral blood mononuclear cells
PBS: phosphate-buffered saline
PCA: principal component analysis
P-gp: P-glycoprotein
PXR: pregnane X receptor
qRT-PCR: quantitative reverse-transcription polymerase chain reaction
RBC: red blood cell
RFU: relative fluorescence units
RIPA: radioimmunoprecipitation assay
RNA-seq: RNA sequencing
RXRα: retinoid X receptor alpha
σ1R: sigma-1 receptor
SD: standard deviation
SNP: single-nucleotide polymorphism
TDF: tenofovir disoproxil fumarate
TFV: tenofovir
TFV-DP: tenofovir diphosphate.

## Acknowledgements

The authors are grateful for ample scholarly discussion with current and past members of the Williams laboratory. We are also grateful for the next generation sequencing services provided by the Emory NBRC Genomics Core (RRID:SCR_026418), supported in part by NIH P51OD011132. Sequencing data was acquired on an Illumina NovaSeq X Plus funded by NIH S10OD038274. Research reported in this publication was supported by the National Institutes of Health under award number R00DA044838, R01DA052859, U01DA058527, and R01DA066140 (DWW), F31DA058562 (ALE), and K00NS118713 (CJW). Additionally, R01DA063076 contributed to a portion of DWW’s effort while this study was ongoing. This research is supported, in part, by the Emory National Primate Research Center Grant No. ORIP/OD P51OD011132, Emory National Primate Research Center Grant No. U42PDP11023, National Institutes of Health’s Office of the Director, Office of Research Infrastructure Programs, National Institutes of Health grant P30AI050409 (Center for AIDS Research at Emory University), and National Institutes of Health grant T32AI157855 (Center for AIDS Research at Emory University). The content is solely the responsibility of the authors and does not necessarily represent the official views of the National Institutes of Health. Graphical images in figures were created using BioRender.

## Data Availability Statement

The authors declare that all the data supporting the findings of this study are available within the paper and its Supplemental Data. The RNA-Seq data discussed in this publication have been deposited in NCBI’s Gene Expression Omnibus (Edgar et al., 2002) and are accessible through GEO Series accession number GSE349670 (https://www.ncbi.nlm.nih.gov/geo/query/acc.cgi?acc=GSE349670).

## Authorship contributions (CRediT statement)

Conceptualization: DWW

Data curation: ASP, RCO, SK, JPC, AE, CJW, JJRF, HNW, AM, KM, TO, PSL, DWW

Formal analysis: ASP, RCO, SK, JPC, AE, CJW, JJRF, HNW, AM, KM, TO, PSL, DWW

Funding acquisition: DWW

Investigation: ASP and DWW

Methodology: ASP, AE, HNW, TO

Supervision: PJG and DWW

Visualization: ASP

Writing – original draft: ASP, AE, DWW

Writing – review and editing: ASP, RCO, SK, JPC, AE, CJW, JJRF, HNW, AM, KM, TO, PSL, KSW, PJG, DWW

## References

1. Zhao, M., et al. Cytochrome P450 Enzymes and Drug Metabolism in Humans. Int. J. Mol. Sci. 22, 12808 (2021).

2. Radke, J., Tocki, K. & Faine, B. A. A Review of the Metabolism and Potential Drug-drug Interactions With Addictive Drugs. J. Addict. Med. 16, e30–e39 (2022).

3. Cho, A. K., Narimatsu, S. & Kumagai, Y. Metabolism of drugs of abuse by cytochromes P450. Addict. Biol. 4, 283–301 (1999).

4. Zhou, S. et al. Interactions of Herbs with Cytochrome P450. Drug Metab. Rev. 35, 35–98 (2003).

5. Hernández-Lorca, M. et al. Dietary Modulation of CYP3A4 and Its Impact on Statins and Antidiabetic Drugs: A Narrative Review. Pharmaceuticals 18, 1351 (2025).

6. Zhou, S.-F. Drugs Behave as Substrates, Inhibitors and Inducers of Human Cytochrome P450 3A4. Curr Drug Metab 9, 310–322 (2008).

7. Zhou, S.-F., Xue, C. C., Yu, X.-Q., Li, C. & Wang, G. Clinically Important Drug Interactions Potentially Involving Mechanism-based Inhibition of Cytochrome P450 3A4 and the Role of Therapeutic Drug Monitoring. Ther. Drug Monit. 29, 687–710 (2007).

8. Lee, J., Beers, J. L., Geffert, R. M. & Jackson, K. D. A Review of CYP-Mediated Drug Interactions: Mechanisms and In Vitro Drug-Drug Interaction Assessment. Biomolecules 14, 99 (2024).

9. Palleria, C. et al. Pharmacokinetic drug-drug interaction and their implication in clinical management. J. Res. Méd. Sci. : Off. J. Isfahan Univ. Méd. Sci. 18, 601–10 (2012).

10. Zang, X. et al. Intracellular islatravir-triphosphate half-life supports extended dosing intervals. Antimicrob. Agents Chemother. 68, e00458–24 (2024).

11. Bazzoli, C. et al. Intracellular Pharmacokinetics of Antiretroviral Drugs in HIV-Infected Patients, and their Correlation with Drug Action. Clin. Pharmacokinet. 49, 17–45 (2010).

12. Jagarapu, A., Piovoso, M. J. & Zurakowski, R. An Integrated Spatial Dynamics—Pharmacokinetic Model Explaining Poor Penetration of Anti-retroviral Drugs in Lymph Nodes. Front. Bioeng. Biotechnol. 8, 667 (2020).

13. Dixit, N. M. & Perelson, A. S. Complex patterns of viral load decay under antiretroviral therapy: influence of pharmacokinetics and intracellular delay. J. Theor. Biol. 226, 95–109 (2004).

14. D’Avolio, A., Pensi, D., Baietto, L. & Perri, G. D. Therapeutic drug monitoring of intracellular anti-infective agents. J. Pharm. Biomed. Anal. 101, 183–193 (2014).

15. Avery, L. B., Zarr, M. A., Bakshi, R. P., Siliciano, R. F. & Hendrix, C. W. Increasing Extracellular Protein Concentration Reduces Intracellular Antiretroviral Drug Concentration and Antiviral Effect. AIDS Res. Hum. Retroviruses 29, 1434–1442 (2013).

16. Lee, J. E. et al. Trends in HIV-Related Services Offered by Substance Abuse Treatment Facilities. Cureus 16, e57400 (2024).

17. Meyer, J. P., Springer, S. A. & Altice, F. L. Substance Abuse, Violence, and HIV in Women: A Literature Review of the Syndemic. J. Women’s Heal. 20, 991–1006 (2011).

18. Gorman, E. M. & Carroll, R. T. Substance Abuse and HIV: Considerations With Regard to Methamphetamines and Other Recreational Drugs for Nursing Practice and Research. J. Assoc. Nurses AIDS Care 11, 51–62 (2000).

19. Angulo-Arreola, I. A., Bastos, F. I. & Strathdee, S. A. Substance Abuse and HIV/AIDS in the Caribbean. J. Int. Assoc. Provid. AIDS Care (JIAPAC) 16, 56–74 (2017).

20. Ortiz, R. C. et al. Cocaine regulates antiretroviral therapy CNS access through pregnane-x receptor-mediated drug transporter and metabolizing enzyme modulation at the blood brain barrier. Fluids Barriers CNS 21, 5 (2024).

21. Wilkins, H. N. et al. All Blood Brain Barrier Cell Types Demonstrate Capability to Influence Differential Tenofovir and Emtricitabine Metabolism and Transport in the Brain. ACS Pharmacol. Transl. Sci. 7, 3626–3640 (2024).

22. Ellison, A. L. et al. Antiviral and anti-inflammatory effects of Cannabidiol in HIV/SIV infection. Brain, Behav., Immun. 106843 (2026) doi:10.1016/j.bbi.2026.106843.

23. Lin, J. & Redies, C. Histological evidence: housekeeping genes beta-actin and GAPDH are of limited value for normalization of gene expression. Dev. Genes Evol. 222, 369–376 (2012).

24. Boujedidi, H. et al. Housekeeping Gene Variability in the Liver of Alcoholic Patients. Alcohol.: Clin. Exp. Res. 36, 258–266 (2012).

25. He, K. et al. Identification of stable housekeeping genes in mouse liver for studying carbon tetrachloride-induced injury and cellular senescence. Sci. Rep. 14, 26544 (2024).

26. Westerberg, L. J. S., Dedic, B., Näslund, E., Thorell, A. & Spalding, K. L. Superior normalization using total protein for western blot analysis of human adipocytes. PLOS One 20, e0328136 (2025).

27. Wilkins, H. N., et al. Drug Metabolism and Transport Capacity of Endothelial Cells, Pericytes, and Astrocytes: Implications for CNS Drug Disposition. bioRxiv 2024.08.01.606165 (2024) doi:10.1101/2024.08.01.606165.

28. Cheng, Q., Sohl, C. D. & Guengerich, F. P. High-throughput fluorescence assay of cytochrome P450 3A4. Nat. Protoc. 4, 1258–1261 (2009).

29. Rosado-Franco, J. J., et al. Roadmap for the expression of canonical and extended endocannabinoid system receptors and metabolic enzymes in peripheral organs of preclinical animal models. Physiol. Rep. 12, e15947 (2024).

30. Williams, D. W., Askew, L. C., Jones, E. & Clements, J. E. CCR2 Signaling Selectively Regulates IFN-α: Role of β-Arrestin 2 in IFNAR1 Internalization. J Immunol 202, 105–118 (2019).

31. Allen, M. I., Johnson, B. N. & Nader, M. A. A comparison of the reinforcing strength of cocaethylene and cocaine in monkeys responding under progressive-ratio and concurrent choice schedules of reinforcement. Drug Alcohol Depend. 251, 110952 (2023).

32. Johnson, B. N., Allen, M. I. & Nader, M. A. Acquisition of cocaine reinforcement using fixed-ratio and concurrent choice schedules in socially housed female and male monkeys. Psychopharmacology 241, 263–274 (2024).

33. Allen, M. I. et al. Behavioral and neuronal extracellular vesicle biomarkers associated with nicotine’s enhancement of the reinforcing strength of cocaine in female and male monkeys. Addict. Neurosci. 11, 100151 (2024).

34. Allen, M. I., Costa, M. B., Johnson, B. N., Gould, R. W. & Nader, M. A. Cognitive performance as a behavioral phenotype associated with cocaine self-administration in female and male socially housed monkeys. Neuropsychopharmacology 1–9 (2024) doi:10.1038/s41386-024-01882-7.

35. John, W. S., Martin, T. J. & Nader, M. A. Cannabinoid Modulation of Food-Cocaine Choice in Male Rhesus Monkeys. J. Pharmacol. Exp. Ther. 373, jpet.119.263707 (2020).

36. Banks, M. L. et al. Effects of Cocaine and MDMA Self-Administration on Serotonin Transporter Availability in Monkeys. Neuropsychopharmacology 33, 219–225 (2008).

37. Nader, M. A. et al. Effects of Cocaine Self-administration on Striatal Dopamine Systems in Rhesus Monkeys: Initial and Chronic Exposure. Neuropsychopharmacology 27, 35–46 (2002).

38. Czoty, P. W. & Nader, M. A. Effects of dopamine D1-like receptor ligands on food-cocaine choice in socially housed male cynomolgus monkeys. J. Pharmacol. Exp. Ther. 379, JPET-AR-2021-000701 (2021).

39. Wakeford, A. G. P. et al. Effects of early life stress on cocaine intake in male and female rhesus macaques. Psychopharmacology 237, 3583–3589 (2020).

40. Nakata, K. et al. Nuclear Receptor-Mediated Transcriptional Regulation in Phase I, II, and III Xenobiotic Metabolizing Systems. Drug Metab. Pharmacokinet. 21, 437 (2007).

41. Jin, P. et al. Nuclear receptors in health and disease: signaling pathways, biological functions and pharmaceutical interventions. Signal Transduct. Target. Ther. 10, 228 (2025).

42. Gee, R. R. F., Huber, A. D. & Chen, T. Regulation of PXR in drug metabolism: chemical and structural perspectives. Expert Opin. Drug Metab. Toxicol. 20, 9–23 (2024).

43. Duc, B. W. L. et al. Norcocaine and N-Hydroxynorcocaine Formation in Human Liver Microsomes: Role of Cytochrome P-450 3A4. Pharmacology 46, 294–300 (1993).

44. Jastrzębska, J. & Daniel, W. A. Cocaine-Induced Time-Dependent Alterations in Cytochrome P450 and Liver Function. Int. J. Mol. Sci. 24, 1632 (2023).

45. Zhang, Y. et al. CYP3A4 and CYP3A5: the crucial roles in clinical drug metabolism and the significant implications of genetic polymorphisms. PeerJ 12, e18636 (2024).

46. Pellinen, P. et al. Kinetic characteristics of norcocaine N-hydroxylation in mouse and human liver microsomes: involvement of CYP enzymes. Arch. Toxicol. 74, 511–520 (2000).

47. Liu, Y. & Matsumoto, R. R. Alterations in Fos-Related Antigen 2 and σ1 Receptor Gene and Protein Expression Are Associated with the Development of Cocaine-Induced Behavioral Sensitization: Time Course and Regional Distribution Studies. J. Pharmacol. Exp. Ther. 327, 187–195 (2008).

48. Narayanan, M., Kulkarni, R., Jiang, S., Kashanchi, F. & Prasad, A. Cocaine augments neuro-inflammation via modulating extracellular vesicle release in HIV-1 infected immune cells. Retrovirology 18, 26 (2021).

49. Sharkey, J., Glen, K. A., Wolfe, S. & Kuhar, M. J. Cocaine binding at σ receptors. Eur. J. Pharmacol. 149, 171–174 (1988).

50. Aguinaga, D. et al. Cocaine Effects on Dopaminergic Transmission Depend on a Balance between Sigma-1 and Sigma-2 Receptor Expression. Front. Mol. Neurosci. 11, 17 (2018).

51. Gekker, G. et al. Cocaine-induced HIV-1 expression in microglia involves sigma-1 receptors and transforming growth factor-β1. Int. Immunopharmacol. 6, 1029–1033 (2006).

52. Niu, F. et al. Cocaine-induced release of CXCL10 from pericytes regulates monocyte transmigration into the CNS. J. Cell Biol. 218, 700–721 (2019).

53. Mai, H. N. et al. Glutathione peroxidase-1 knockout potentiates behavioral sensitization induced by cocaine in mice via σ-1 receptor-mediated ERK signaling: A comparison with the case of glutathione peroxidase-1 overexpressing transgenic mice. Brain Res. Bull. 164, 107–120 (2020).

54. Romieu, P., Phan, V.-L., Martin-Fardon, R. & Maurice, T. Involvement of the Sigma1 Receptor in Cocaine-induced Conditioned Place Preference: Possible Dependence on Dopamine Uptake Blockade. Neuropsychopharmacology 26, 444–455 (2002).

55. Romieu, P., Martin-Fardon, R. & Maurice, T. Involvement of the σ1 receptor in the cocaine-induced conditioned place preference. NeuroReport 11, 2885–2888 (2000).

56. Yao, H. et al. Molecular mechanisms involving sigma receptor–mediated induction of MCP-1: implication for increased monocyte transmigration. Blood 115, 4951–4962 (2010).

57. Su, T.-P. Non-canonical Targets Mediating the Action of Drugs of Abuse: Cocaine at the Sigma-1 Receptor as an Example. Front. Neurosci. 13, 761 (2019).

58. Mateu-Mollá, J., Pérez-Gálvez, B. & Villanueva-Blasco, V. J. Pharmacological treatment for substance use disorder: A systematic review. Addict. Behav. 163, 108242 (2025).

59. Knowles, L. G., Armanious, A. J., Peng, Y., Welsh, W. J. & James, M. H. Recent advances in drug discovery efforts targeting the sigma 1 receptor system: Implications for novel medications designed to reduce excessive drug and food seeking. Addict. Neurosci. 8, 100126 (2023).

60. Menkel, M., Terry, P., Pontecorvo, M., Katz, J. L. & Witkin, J. M. Selective σ ligands block stimulant effects of cocaine. Eur. J. Pharmacol. 201, 251–252 (1991).

61. Narayanan, S., Mesangeau, C., Poupaert, J. H. & McCurdy, C. R. Sigma Receptors and Cocaine Abuse. Curr. Top. Med. Chem. 11, 1128–1150 (2011).

62. López, O. V. et al. Sigma-1 Receptor Antagonist (BD1047) Decreases Cathepsin B Secretion in HIV-Infected Macrophages Exposed to Cocaine. J. Neuroimmune Pharmacol. 14, 226–240 (2019).

63. Tsai, S.-Y. A. et al. Sigma-1 receptor mediates cocaine-induced transcriptional regulation by recruiting chromatin-remodeling factors at the nuclear envelope. Proc. Natl. Acad. Sci. 112, E6562–E6570 (2015).

64. Aguinaga, D. et al. The sigma-1 receptor as key common factor in cocaine and food seeking behaviors. J. Mol. Endocrinol. 1, R81–R92 (2019).

65. Gavegnano, C. et al. Cellular Pharmacology and Potency of HIV-1 Nucleoside Analogs in Primary Human Macrophages. Antimicrob. Agents Chemother. 57, 1262–1269 (2012).

66. Hoque, T. M. D. et al. Antiretroviral Drug Transporters and Metabolic Enzymes in Circulating Monocytes and Monocyte-Derived Macrophages of ART-Treated People Living With HIV and HIV-Uninfected Individuals. JAIDS J. Acquir. Immune Defic. Syndr. 87, 1093–1101 (2021).

67. Dhillon, N. K. et al. Cocaine-mediated enhancement of virus replication in macrophages: implications for human immunodeficiency virus-associated dementia. J Neurovirol 13, 483–95 (2007).

68. Wheeler, A. M. et al. Achieving a Deeper Understanding of Drug Metabolism and Responses Using Single-Cell Technologies. Drug Metab. Dispos. 51, 350–359 (2023).

69. Pirmohamed, M. Drug metabolism. Medicine 36, 355–359 (2008).

70. Jhajra, S. et al. Encyclopedia of Drug Metabolism and Interactions. Encycl. Drug Metab. Interact. 1–99 (2013) doi:10.1002/9780470921920.edm028.

71. Ali, H. et al. The impact of the exposome on cytochrome P450-mediated drug metabolism. Front. Pharmacol. 16, 1639646 (2025).

72. Richert, L. et al. Use of mRNA expression to detect the induction of drug metabolising enzymes in rat and human hepatocytes. Toxicol. Appl. Pharmacol. 235, 86–96 (2009).

73. Ekiciler, A. et al. Quantitative Cytochrome P450 3A4 Induction Risk Assessment Using Human Hepatocytes Complemented with Pregnane X Receptor-Activating Profiles. Drug Metab. Dispos. 51, 276–284 (2023).

74. Mattinen, L., Kublbeck, J., Rechardt, O., Honkakoski, P. & Rautio, J. Direct and Rapid Transcript Analysis Assay for CYP mRNA Expression and Inducibility in Human Primary Hepatocytes. Drug Metab. Lett. 8, 77–87 (2015).

75. Fahmi, O. A., Kish, M., Boldt, S. & Obach, R. S. Cytochrome P450 3A4 mRNA Is a More Reliable Marker than CYP3A4 Activity for Detecting Pregnane X Receptor-Activated Induction of Drug-Metabolizing Enzymes. Drug Metab. Dispos. 38, 1605–1611 (2010).

76. Li, D., Gaedigk, R., Hart, S. N., Leeder, J. S. & Zhong, X. The Role of CYP3A4 mRNA Transcript with Shortened 3′-Untranslated Region in Hepatocyte Differentiation, Liver Development, and Response to Drug Induction. Mol. Pharmacol. 81, 86–96 (2012).

77. Okawa, M. et al. Circadian proteolysis by the E3 ubiquitin ligase gp78 controls CYP3A4 activity in human hepatocytes. Biochem. Pharmacol. 253, 118330 (2026).

78. Aquaro, S., Borrajo, A., Pellegrino, M. & Svicher, V. Mechanisms underlying of antiretroviral drugs in different cellular reservoirs with a focus on macrophages. Virulence 11, 400–413 (2020).

79. Gavegnano, C. & Schinazi, R. F. Antiretroviral Therapy in Macrophages: Implication for HIV Eradication. Antivir Chem Chemother 20, 63–78 (2009).

80. Herskovitz, J. & Gendelman, H. E. HIV and the Macrophage: From Cell Reservoirs to Drug Delivery to Viral Eradication. J Neuroimmune Pharm 14, 52–67 (2018).

81. Joseph, J. et al. Role of macrophages in HIV pathogenesis and cure: NIH perspectives. J. Leukoc. Biol. 112, 1233–1243 (2022).

82. He, S., Zubair, A. & Al-Emam, A. Tissue Reservoirs of HIV: The Hidden Protective Role of Lymphoid and Myeloid Cells. Rev. Méd. Virol. 36, e70159 (2026).

83. Corsi, F. et al. Antiretroviral Therapy through Barriers: A Prominent Role for Nanotechnology in HIV-1 Eradication from Sanctuaries. J. Pharm. Pharmacol. 4, (2016).

84. Devanathan, A. S. & Cottrell, M. L. Pharmacology of HIV Cure: Site of Action. Clin. Pharmacol. Ther. 109, 841–855 (2021).

