## Supplemental Figures for "Primary Human Macrophages Exhibit Functional CYP3A4 Activity with Marked Expression-Function Discordance and Cocaine-Induced Catalytic Activation"

**Supplemental Data**
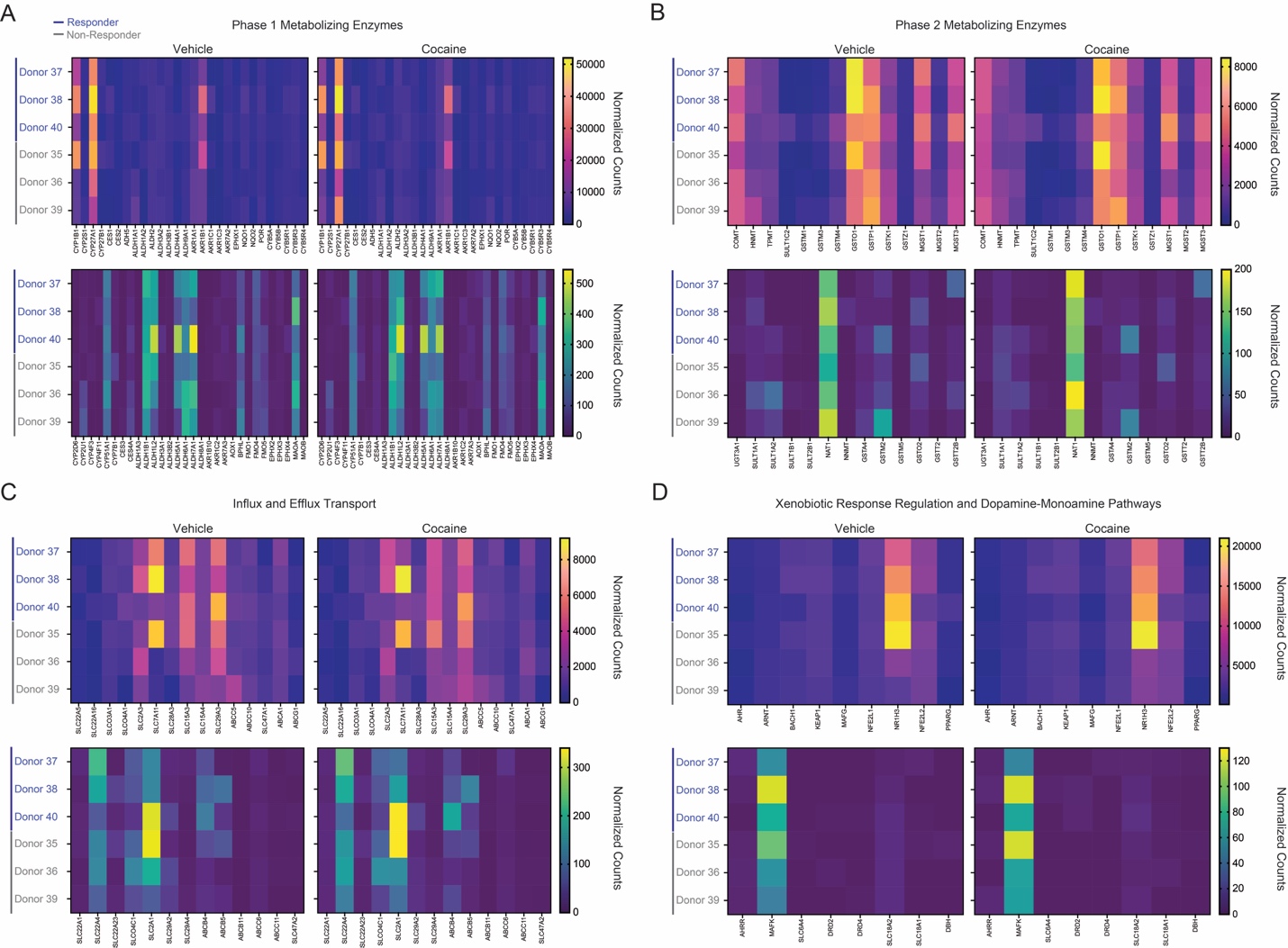

**Supplemental Figure 1: Donor-dependent transcriptional responses of selected drug metabolism, transport, and xenobiotic response genes following MDM cocaine treatment**. Heatmaps depict normalized transcript counts across six donors following vehicle or cocaine treatment. Donors are shown in the same order across panels and classified according to their CYP3A4 response profile as “Responders” (blue; Donors 37, 38, and 40) or “Non-responders” (gray; Donors 35, 36, and 39). Genes were grouped by expression abundance for visualization: (A) Phase I metabolizing enzymes, (B) Phase II metabolizing enzymes, (C) influx and efflux transport, and (D) xenobiotic response regulation and dopamine–monoamine pathways. Color scales indicate normalized transcript counts and were independently scaled for each expression-abundance group. Color intensity represents normalized counts according to the scale shown adjacent to each heatmap.
